# The high-resolution *in vivo* measurement of replication fork velocity and pausing by lag-time analysis

**DOI:** 10.1101/2022.08.13.503874

**Authors:** Dean Huang, Anna E. Johnson, Brandon S. Sim, Teresa Lo, Houra Merrikh, Paul A. Wiggins

**Affiliations:** Department of Physics, University of Washington, Seattle, Washington 98195, USA; Department of Biochemistry, Vanderbilt University, Nashville, Tennessee 37205, USA; Department of Pathology, Microbiology, and Immunology, Vanderbilt University Medical Center, Nashville, Tennessee 37232, USA; Department of Bioengineering, University of Washington, Seattle, Washington 98195, USA; Department of Microbiology, University of Washington, Seattle, Washington 98195, USA

## Abstract

An important step towards understanding the mechanistic basis of the central dogma is the quantitative characterization of the dynamics of nucleic-acid-bound molecular motors in the context of the living cell, where a crowded cytoplasm as well as competing and potentially antagonistic processes may significantly affect their rapidity and reliability. To capture these dynamics, we develop a novel method, lag-time analysis, for measuring *in vivo* dynamics. The approach uses exponential growth as the stopwatch to resolve dynamics in an asynchronous culture and therefore circumvents the difficulties and potential artifacts associated with synchronization or fluorescent labeling. Although lag-time analysis has the potential to be widely applicable to the quantitative analysis of *in vivo* dynamics, we focus on an important application: characterizing replication dynamics. To benchmark the approach, we analyze replication dynamics in three different species and a collection of mutants. We provide the first quantitative locus-specific measurements of fork velocity, in units of kb per second, as well as replisome-pause durations, some with the precision of seconds. The measured fork velocity is observed to be both locus and time dependent, even in wild-type cells. In addition to quantitatively characterizing known phenomena, we detect brief, locus-specific pauses at rDNA in wild-type cells for the first time. We also observe temporal fork velocity oscillations in three highly-divergent bacterial species. Lag-time analysis not only has great potential to offer new insights into replication, as demonstrated in the paper, but also has potential to provide quantitative insights into other important processes.

## I. INTRODUCTION

At a single-molecule scale, all cellular processes are both highly stochastic as well as subject to a crowded cellular environment where they typically compete with a large number of potentially-antagonistic processes that share the same substrate [1, 2]. In spite of these challenges, essential processes must be robust at a cellular scale to facilitate efficient cellular proliferation. Understanding how these processes are regulated to achieve robustness remains an important and outstanding biological question [3–9]. However, a central challenge in investigating these questions is the quantitative characterization of the activity of enzymes in the context of the living cell. For instance, although single-molecule assays can resolve the pausing of molecular motors on nucleic-acid substrates in the context of *in vitro* measurements [10, 11], performing analogous measurements in the physiologically-relevant environment of the cell, where these processes are subject to antagonism, poses a severe challenge to the existing methodologies [12].

In this paper, we develop an approach, *lag-time analysis*, that facilitates the quantitative characterization of dynamics, with resolution of seconds, in the context of the living cell. The approach exploits exponential growth as the stopwatch to capture dynamics in exponentially proliferating cellular cultures [13] and unlike competing approaches, it can circumvent the difficulties and potential artifacts introduced by cell synchronization [14] or fluorescent labeling. Although in principle the approach is broadly applicable to all central dogma processes (*i*.*e*. replication, transcription, and translation) as well as other dynamics, we will focus on replication dynamics exclusively for concreteness.

In this paper, we demonstrate the first locus-specific genome-wide measurement of the fork velocity, in units of kilobases per second, and the duration of replisome pauses. It facilitates detailed comparisons to be made, not just between different loci in a single cell, but between wild-type and mutant cells as well as between bacterial species. We apply this approach to analyze three model bacterial systems: *Bacillus subtilis, Vibrio cholerae*, and *Escherichia coli*. In *B. subtilis*, we analyze transcription-induced replication antagonism which is the main determinant of replisome dynamics in a set of mutants with retrograde (reverse-oriented) fork motion. An analysis of *V. cholerae* provides evidence that fork number is an important determinant of fork velocity, but also provides clear evidence that fork velocity is time dependent. To explore this time-dependence, we analyze the fork-velocity in *E. coli* which provides strong evidence for temporally-oscillating fork velocity, consistent with a recent report [15]. Finally, we demonstrate that these oscillations are observed in all three organisms. In summary, the observed phenomena demonstrate the central importance of characterizing central dogma processes in the context of the living cell, where their activity is regulated and modulated by the cellular environment.

## II. RESULTS

### The bacterial cell cycle

The bacterial cell cycle is divided into three periods [16, 17]: The B period is analogous to the G_1_ phase of the eukaryotic cell cycle, corresponding to the period between cell birth and replication initiation. The C period is analogous to the S phase (and early M phase) in which the genome is replicated and simultaneously and sequentially segregated [18]. The D period is analogous to a combination of phases G_2_ and late-M, corresponding to a period of time between replication termination and cell division, including the process of septation (*i*.*e*. cytokinesis).

The demographics of cell-cycle periods of exponentially-growing bacterial cells were first quantitatively modeled by Cooper and Helmstetter in an influential paper [19] and then refined by multiple authors [20–22]. In the Methods Section, we generalize these models to demonstrate that the measured marker-frequency analysis quantitatively measures the cell-cycle replication dynamics. The key results are summarized below.

### Lag-time analysis

Our strategy will be *to use exponential growth as the stopwatch with which we resolve cell-cycle dynamics*. In short, cells with greater cell-cycle progression (*i*.*e*. age) are depleted in the population, equivalent to an independent, exponentially-proliferating species that *lags* newborn cells by a time equal to its age [13]. (See Fig. 1 for a schematic illustration of the approach.) *Lag-time analysis* is the measurement of this time lag. In principle, this approach can be applied to characterize the dynamics of any biological molecules or complexes; however, for concreteness, we will focus on replication dynamics since the replication process is of great biological interest and next-generation sequencing provides a powerful tool for digital, as well as genome-wide, quantitation of the number of genomic loci.

**FIG. 1.**
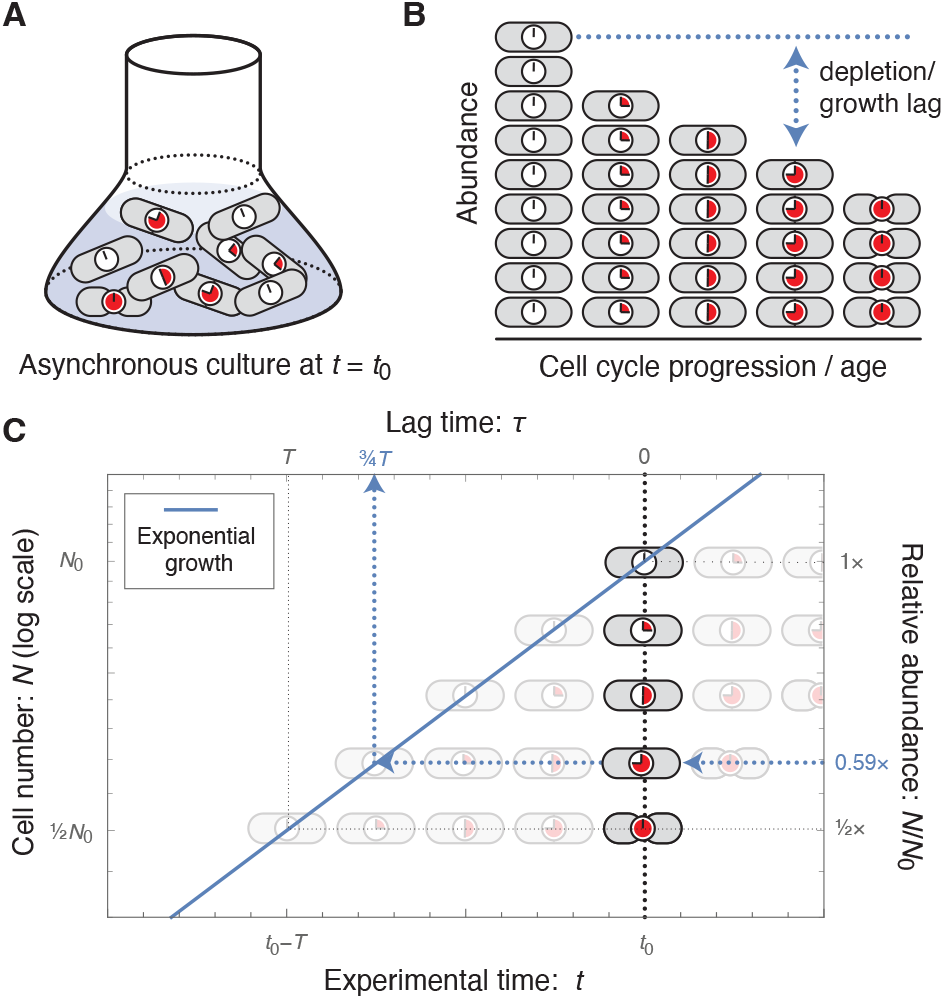
Lag-time analysis. Panel A: Sample preparation. An asynchronous culture in steady-state exponential growth is harvested at time *t* = *t*_0_. **Panel B: Quantitation of demographics**. Cell abundance is quantified. For analyzing replication dynamics, cell quantitation is performed by nextgeneration sequencing. **Panel C: Measurement of lag time**. The dotted black line represents the culture at *t* = *t*_0_. Cells with greater cell cycle progression (*i*.*e*. age) are depleted relative to newborn cells. For each cell age, the relative abundance determines the *lag time*. Their abundance is equivalent to an exponentially proliferating species that lags newborn cells by a time equal to its age. For instance, the nine-o’clock cell is at a relative abundance of 0.59 with a lag time of 3/4ths the mass-doubling time *T*. Schematically, start from the observed number of nine-o’clock cells, and follow that lineage horizontally (back in time) until you reach the newborn cell, born at *t* = *t*_0_ −*τ* (blue dotted line). For a stochastic cell cycle, lag time measures the exponential mean of the stochastic time (Eq. 2).

In marker-frequency experiments, the number of each sequence *N* (𝓁) in a steady-state, asynchronously growing population is determined by mapping next-generation-sequencing reads to the reference genome. This marker frequency can be reinterpreted as a measurement of the *lag time τ* (𝓁):

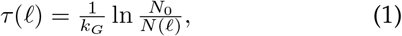

where *N* (𝓁) is the observed number of the locus at genomic position 𝓁 and *N*_0_ is the observed number of the origin in the culture and *k*_*G*_ is the growth rate. This relation can be understood as a consequence of the exponential growth law [13].

In a deterministically-timed model, the measured lag time would be equal to the replication time relative to initiation. In reality, the timing of all processes in the cell cycle is stochastic. We previously showed that the measured lag time is related to the distribution of durations in single cells by the exponential mean [13]:

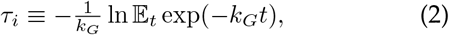

where 𝔼_*t*_ is the expectation over stochastic time *t* with distribution *t* ∼ *p*_*i*_(·).

### Determination of replisome pause durations

Replisome pause durations or the lag time difference between the replication of any two loci can be computed using the difference of lag times between the two loci:

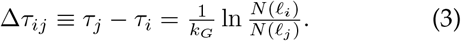

We emphasize that the observed lag-time difference is the exponential mean of the stochastic time difference, which has important consequences for slow processes.

### Determination of the fork velocity

For fast processes, like single nucleotide incorporation, the exponential mean leads to a negligible correction (see Methods); therefore, the fork velocity has a simple interpretation: it is the slope of the genomic position versus lag-time curve:

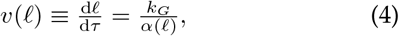

or equivalently it is the ratio of the growth rate to the log-slope:

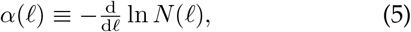

which can be directly determined from the marker frequency.

### Lag-time analysis reveals *V. cholerae* replication dynamics

To explore the application of lag-time analysis to characterize replication dynamics, we begin our analysis in the bacterial model system *Vibrio cholerae*, which harbors two chromosomes: Chromosome 1 (Chr1) is 2.9 Mb and Chromosome 2 (Chr2) is 1.1 Mb. The origin of Chr1, *oriC1*, fires first and roughly the first half of replication is completed before the replication-initiator-RctB-binding-site *crtS* is replicated, triggering Chr2 initiation at *oriC2* [23–25]. Chr1 and Chr2 then replicate concurrently for the rest of the C period. (See Fig. 2A.)

**FIG. 2.**
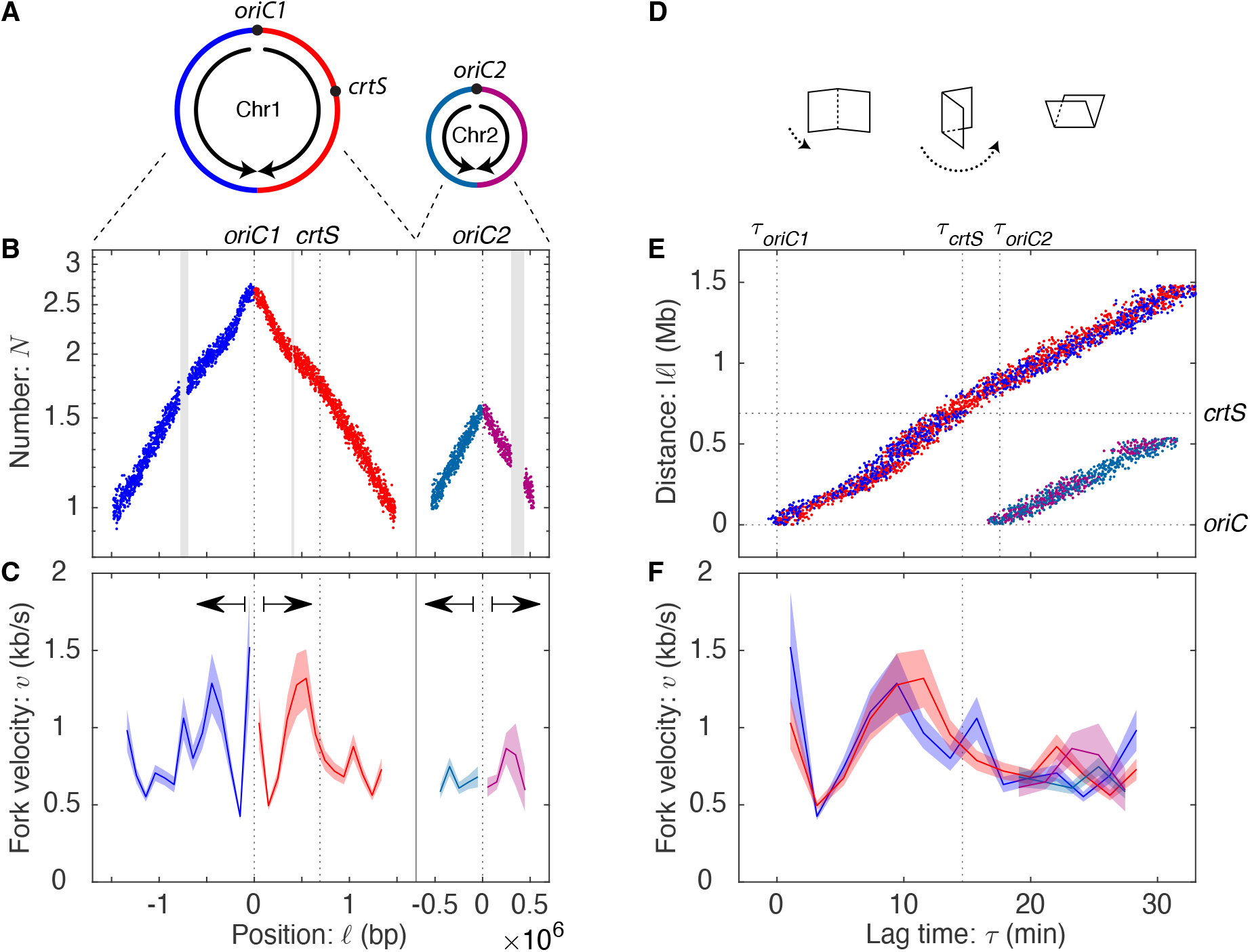
Panel A: Chromosome organization in *V. cholerae*. *V. cholerae* harbors two chromosomes Chr1 and Chr2. *oriC2* initiates shortly after the *crtS* sequence is replicated on the right arm of Chr1. **Panel B: Marker frequency for *V. cholerae* grown on LB**. Repetitive sequences that cannot be mapped result in gaps. **Panel C: Fork velocity is locus-dependent**. The fork velocity is shown as a function of genomic position with an error region. Statistically significant differences in the fork velocity are observed between loci. There is significant bilateral (*i*.*e*. mirror) symmetry around the origin. **Panel D:** A visual representation of the relation between the log-marker-frequency and lag-time plots: Fold at the origin and rotate. **Panel E: Lag-time analysis**. The replication forks start at the origin at lag time zero and then accelerate and decelerate synchronously, as the forks move away from the origin. The consistency in arm position is a manifestation of bilateral symmetry. **Panel F: Fork velocity as a function of lag time**. In addition to bilateral symmetry, after Chr2 initiates, all four forks show roughly consistent velocities.

To demonstrate the power of lag-time analysis, we compute the marker frequency, lag-time, and fork velocities. To measure pause times and replication velocities, we generate a piecewise linear model with a resolution set by the Akaike Information Criterion (AIC). The AIC-optimal model for fast growth (in LB) had 39 knots, spaced by 100 kb, generating 38 measurements of locus velocity across the two chromosomes. The replication dynamics for growth in LB is shown in Fig. 2.

**TABLE I.** Fork number and velocities under different growth conditions. Increasing fork number by increasing cell metabolism does not consistently reduce fork velocity. Fork velocities in fast growth are higher in *E. coli* and lower in *V. cholerae*. The statistical significance column shows the p-value for the null hypothesis of constant fork velocity. For more details on how these values are calculated, see Supplemental Material Sec. IV.

| Organism | Growth condition | Doubling time: $T$ (min) | Fork statistics | | | | | Statistical significance p-value: $p$ |
| --- | --- | --- | --- | --- | --- | --- | --- | --- |
| | | | C period: $C$ (min) | Fork number: $\bar{N}_F$ | Velocity mean: $\bar{v}$ (kb/s) | std: $\sigma_v$ (kb/s) | Symmetry: $f_S$ | |
| *E. coli* | LB | 19 | 30 | 3.8 | 1.3 | 0.19 | 84% | $\ll 10^{-30}$ |
| | M9 | 69 | 46 | 1.2 | 0.85 | 0.12 | 59% | $6 \times 10^{-12}$ |
| *V. cholerae* | LB | 22 | 31 | 4.3 | 0.82 | 0.27 | 76% | $\ll 10^{-30}$ |
| | M9 | 50 | 32 | 1.5 | 0.84 | 0.28 | 70% | $\ll 10^{-30}$ |
| *B. subtilis*<br>*rrnIHG* | S7 | 64 | 42 | 0.82 | 1.1 | 0.68 | 50% | $\ll 10^{-30}$ |
| | MOPS+CA | 44 | 40 | 1.2 | 0.86 | 0.45 | 45% | $\ll 10^{-30}$ |
| | MOPS | 50 | 41 | 1.1 | 0.99 | 0.52 | 57% | $\ll 10^{-30}$ |

### Lag-time analysis enables the measurement of the duration of fast processes

We focus first on the duration of time between *crtS* replication and the initiation of *oriC2*. Fluorescence microscopy imaging reveals that this wait time is very short [26], but it is very difficult to quantify since the precise timing of the replication of the *crtS* sequence is difficult to determine by fluorescence imaging; however, this is a natural application for lag-time analysis. To measure the lag-time difference between *crtS* replication and *oriC2* replication, we use Eq. 3 to compute the replication time difference from the relative copy numbers. For this analysis, we generate a piecewise linear model with knots at the *crtS* and *oriC2* loci. The measured lag time is

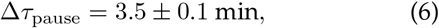

a pause duration which is clearly resolved in the lag-time plot shown in Fig. 2E.

### The fork velocity is locus dependent

It is qualitatively clear from the fork-distance-versus-time plot (Fig. 2E) that the fork velocity is locus dependent since the trajectory is not straight. To test this question statistically, we compare the 39-knot model to the null hypothesis (constant fork velocity), which is rejected with a p-value of *p* ≪ 10^−30^ and therefore the data cannot be described by a single fork velocity. (See Tab. I.) The resulting velocity profiles are shown in Fig. 2CF.

### Bilateral symmetry supports a time-dependent mechanism

Our understanding of the replication process motivated two general classes of mechanisms: (i) *time-dependent* and (ii) *locus-dependent* mechanisms. Time-dependent mechanisms, like a dNTP-limited replication rate, affect all forks uniformly and therefore loci equidistant from the origin should have identical fork velocities:

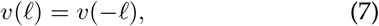

where 𝓁 is the genetic position relative to the origin. In contrast, in a locus-dependent mechanism, like replication-conflict-induced slowdowns, the slow regions are expected to be randomly distributed over the chromosome. In this scenario we expect to see no bilateral symmetry between arms (*e*.*g*. Fig. 7B).

A bilateral symmetry between the arms is clearly evident in the data (the mirror symmetry about the origin in Figs. 2BC and is manifest in the lag-time analysis as the coincidence between the left and right arm trajectories and velocities in Figs. 2EF. To quantitate this symmetry, we divide the variance of the fork velocity into symmetric and antisymmetric contributions. (See the Supplemental Material Sec. IV A.) A time-dependent mechanism would generate a *f*_*S*_ = 100% symmetric variance, whereas a locus-dependent mechanism would be expected to generate equal symmetric and antisymmetric variance contributions (*f*_*S*_ = 50%). *V. cholerae* Chr1 and Chr2 have *f*_*S*_ = 76% symmetry, consistent with a time-dependent mechanism playing a dominant but not exclusive role in determining the fork velocity. (See Tab. I.)

### The replisome pauses briefly at rDNA in *B. subtilis*

To explore the possibility that locus-dependent mechanisms could play a dominant role in determining the fork velocity profile, we next characterized the fork dynamics in the context of replication conflicts, where the antagonism between active transcription and replication, have been reported to stall the replisome by a locus-specific mechanism [9, 27]. In *B. subtilis*, there are seven highly-transcribed rDNA loci on the right arm and only a single locus of the left arm. Consistent with the notion of rDNA-induced pausing, the *ter* locus is positioned asymmetrically on the genome, at 172° rather than 180°, leading the right arm of the chromosome to be shorter than the left arm. (See Fig. 3A.) In spite of the difference in length, both arms terminate roughly synchronously, implying that the average fork velocity is lower on the right arm, consistent with putative fork pausing at the rDNA loci. Are these conflict-induced pauses present in wild-type cells where the replication and transcription are co-directional? We have previously reported evidence based on single-molecule imaging that they are [12], but there is as of yet no other unambiguous supporting evidence.

**FIG. 3.**
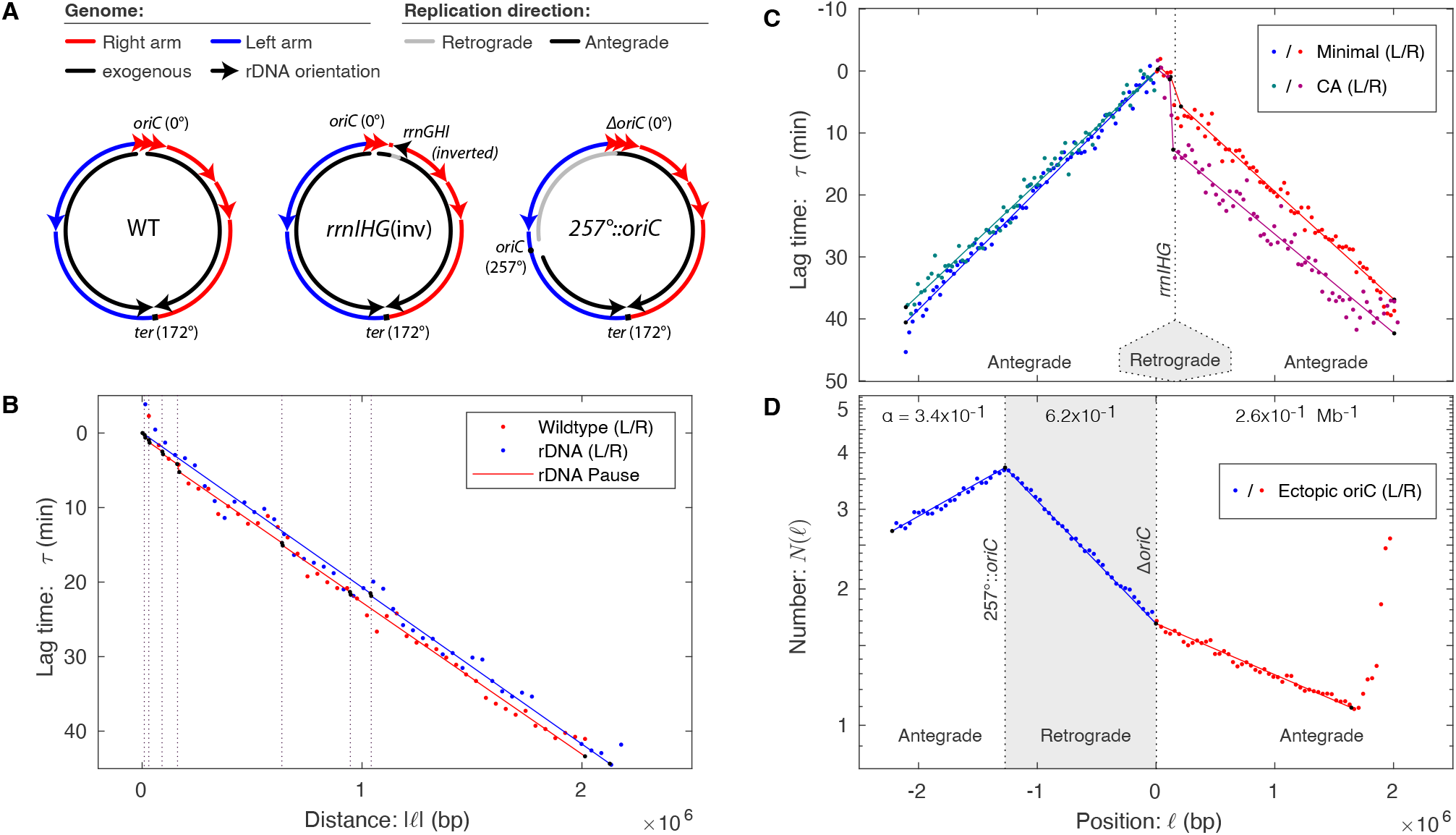
*B. subtilis* fork dynamics and transcriptional conflicts. Panel A: Chromosomal structure for wild-type and mutant *B. subtilis* strains. The *ter* region in wild-type *B. subtilis* is positioned at 172°, rather than 180°, making the right arm shorter than the left. In *rrnIHG*(inv), the *rrnIHG* locus is inverted so that it is transcribed in a head-on orientation with respect to replication. In 257°::*oriC*, the origin is moved to 257°, resulting in a short left arm that terminates at the terminus and a long right arm that replicates initially in the retrograde direction, before replicating the residuum of the right arm in the antegrade orientation. **Panel B: Lag time in wild-type cells**. Replication on the right arm (red) is delayed relative to the left arm (blue) by multiple endogenous co-directional rDNA loci. **Panel C: Head-on conflicts lead to pausing**. The *rrnIHG* genes are inverted so that transcription of the rDNA locus is in the head-on direction. A longer lag-time pause is observed at intermediate growth rates (CA, purple) than slow growth (minimal, red). Fork velocities elsewhere are roughly consistent. **Panel D: Retrograde fork motion is slow**. The retrograde fork motion in R is slow compared to antegrade replication in A1. Late antegrade motion in A2 is faster than early antegrade motion in A1.

To detect putative short pauses at the rDNA loci in wild-type *B. subtilis*, a low-noise dataset was essential. We therefore examined a number of different datasets, including our own, to search for a dataset with the lowest statistical and systematic noise. A marker-frequency dataset for a nearly wild-type strain growing on minimal media was identified for which the noise level was extremely low. (See Supplemental Material Sec. III C 2.) The lag-time analysis is shown in Fig. 3B. Replication pauses should result in discrete steps in the lag time (*e*.*g*. Fig. 7B); however, no clearly-defined steps are visible in the lag-time plot. The pauses are either absent or too small to be clearly visible without statistical analysis.

To achieve optimal statistical resolution, we used the AIC model-selection framework [28, 29] on four competing hypotheses: In Model 1, fork velocities are constant and equal on both arms with no pauses. In Model 2, fork velocities are constant but unequal on the left and right arms with no pauses. In Model 3, fork velocities are constant and equal on the left and right arms with equal-duration pauses at each rDNA locus. In Model 4, fork velocities are constant and unequal on the left and right arms with equal-duration pauses at each rDNA locus. AIC selected Model 3 (equal arm velocities with rDNA pauses) and a pause duration of:

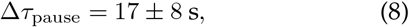

is observed. The pause models were strongly supported over the non-pause models (ΔAIC_23_ = 4.3 and ΔAIC_43_ = 9.4). Therefore, statistical analysis supports the existence of short slowdowns (*i*.*e*. pauses) at the rDNA, even if these features are not directly observable without statistical analysis. In higher-noise datasets, the statistical inference was ambiguous.

### Strong, head-on conflicts lead to long pauses

Although we have just demonstrated that endogenous co-directional conflicts are detected statistically, they do not lead to a clear unambiguous signature. In contrast, strong, exogenous head-on conflicts in which the replisome and transcriptional machinery move in opposite directions can lead to particularly potent conflicts and even cell death [3–9, 30]. The ability to engineer conflicts at specific loci facilitates the use of lag-time analysis for measuring the duration of the replication pauses.

To measure the pause durations due to head-on conflicts, we analyze the marker frequency from a strain, *rrnIHG*(inv), generated by Srivatsan and coworkers with three rDNA genes (*rrnIHG*) inverted so that they are transcribed in the head-on orientation. Marker-frequency datasets were reported for this strain in two growth conditions: minimal supplemented with casamino acids, in which the strain grows at an intermediate growth rate, and unsupplemented minimal media [31]. (Mutant cells cannot proliferate in rich media, presumably because the transcription conflicts are so severe [31].) In both slow and intermediate growth conditions, a clearly resolved step at the head-on locus is observed in the marker-frequency and lag-time analysis (Fig. 3B), exactly analogous to the simulated pause (Fig. 7B).

To determine the pause durations in the two growth conditions, we again consider a model with an unknown pause duration (at the inverted rDNA locus) and constant but unequal fork velocities on the left and right arms. The observed lag-time pauses are

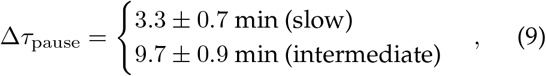

for the slow and intermediate growth rates, respectively. Although lag-time analysis reports a precise pause duration, it is important to remember that the observed lag time corresponds to the exponential mean of the stochastic state lifetime (Eq. 2), including cells that arrest and therefore never complete the replication process. Eq. 18 accounts for the pause generated by this arrested cell fraction. Srivatsan and coworkers report that 10% of the cells are arrested in intermediate growth, which accounts for 8.3 min of the lag time, leaving an estimated pause time of Δ*τ*_pause_ = 1.4 ± 0.9 min for non-arrested cells, which is roughly consistent with the pause time observed in slow growth conditions.

### Slow retrograde replication in *B. subtilis*

Are all conflict-induced slowdowns consistent with long pauses at a small number of rDNA loci? Wang et al. have previously engineered a head-on strain, 257°::*oriC*, with less severe conflicts by moving *oriC* down the left arm of the chromosome to 257° [32]. (See Fig. 3A.) The resulting strain has a very short left arm and a very long right arm, the first third of which is replicated in the *retrograde* (*i*.*e*. reverse to wild-type) orientation. This retrograde region contains only a single rDNA locus. All other regions are replicated in the *antegrade* (*i*.*e*. endogenous) orientation.

Consistent with the analysis of Wang et al., we position knots to divide the chromosome into three regions with three distinct slopes: an early antegrade region *A*1 (the short left arm) with log-slope *α*_*A*1_ = 0.34 ± 0.01 Mb^−1^, a retrograde region *R* with log-slope *α*_*R*_ = 0.63 ± 0.01 Mb^−1^ and a late antegrade region *A*2 with log-slope *α*_*A*2_ = 0.26 ± 0.01 Mb^−1^, that replicates after the left arm terminates. (See Fig. 3C.) Due to the higher percentage of head-on genes in the *R* region compared with the *A*1 region, the conflict model predicts more rapid replication in region *A*1 versus *R*. Consistent with this prediction, the ratio of replication velocities is:

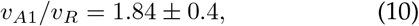

revealing a strong replication-direction dependence. The slope appears relatively constant, consistent with a model of uniformly-distributed slow regions rather than a small number of long pauses as observed in the reversal of the rDNA locus *rrnIHG*. Our quantitative analysis is consistent with the interpretation of Wang et al. [32].

### Rapid late replication due to genomic asymmetry

This dataset has a striking feature that is not emphasized in previous reports: Late antegrade fork velocity is faster than early antegrade velocity:

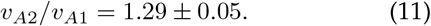

Although this effect is weaker than the replication-direction dependence discussed above (Eq. 10), its size is still comparable. An analogous late-time speedup is seen in two other ectopic origin strains. (See the Supplemental Material Secs. VI H and VI J.)

One potential hypothesis is that a locus-dependent mechanism slows the fork in the *A*1 region relative to the *A*2 region; however, no velocity difference is evident in these regions in the wild-type cells (Fig. 3B). Alternatively, one could hypothesize that there is some form of communication between forks that leads to a slowdown in region *A*1 due to the slowdown in region *R*; however, no coincident slowdown is observed in *rrnIHG*(inv) at a position opposite the *rrnIHG* locus, inconsistent with this hypothesis. Another possible hypothesis is that late-time replication is always rapid; however, no significant speedup is observed in either wild-type *B. subtilis* (Fig. 3A) or *V. cholerae* cells at the end of the replication process (Fig. 3B and Fig. 2E). However, there is one extremely important difference between 257°::*oriC* and the wild-type strains: Due to the asymmetric positioning of the origin and replication traps at the terminus (Fig. 3A), there is only a single active replication fork as the A2 region is replicated. We therefore hypothesize that the fork velocity is inversely related to active fork number.

### Fork number determines velocities in *V. cholera*

To explore the effects of changes in the fork number on fork velocity, it is convenient to return to *V. cholerae*. In slow growth conditions, the cells start the C period with a pair of replication forks, for which the fork-number model predicts faster fork velocity, and finish the replication cycle with two pairs of forks, predicting slower fork velocity.

Although the structure of the velocity profile is more complex than predicted by the fork-number model alone, the observed fork velocity is broadly consistent with its predictions. If a mean fork velocity is computed before and after *oriC2* initiates, the ratio is:

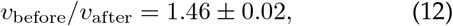

which is quantitatively consistent with the hypothesis that more forks lead to a slowdown in replication and the size of the effect is comparable to what is observed in *B. subtilis* (Eq. 11).

A mutant *V. cholerae* strain has been constructed that facilitates a non-trivial test of the fork-number model: In the monochromosomal strain MCH1, Chr2 is recombined into Chr1 at the terminus of Chr1, resulting in a single monochromosome (Chr 1-2). (See Fig. 4A.) Both the wild-type and MCH1 strains have essentially identical sequence content, implying the locus-dependent model would predict identical replication velocities; however, all replication in MCH1 occurs with only a single set of forks whereas the wild-type strain replicates the latter half of the C period with two pairs of forks, one pair on each chromosome.

**FIG. 4.**
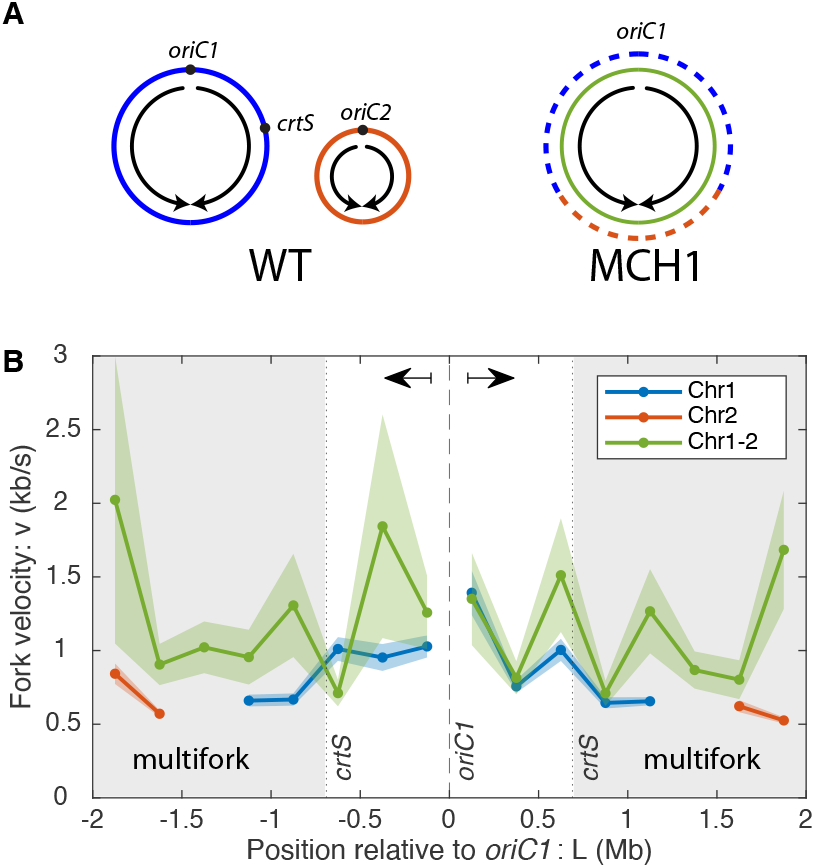
Reducing fork number increases fork velocity. Panel A: The monochromosomal strain MCH1 has a single chromosome (green) which was constructed by recombining Chr2 (orange) into the terminus of Chr1 (blue) [33]. Under slow growth conditions the first part of the chromosome in both strains is replicated by a single pair of forks. When the fork reaches the *crtS* sequence on the right arm, Chr2 is initiated at *oriC2* in the wild-type cells. All of Chr2 and the residuum of Chr1 replicate simultaneously, resulting in two pairs of active forks. In contrast, all sequences in MCH1 are replicated using a single pair of forks. **Panel B:** In MCH1, where all sequences are replicated by a single pair of forks, the fork velocity is faster than is observed in WT cells during the multifork region (grey– sequences replicated after *crtS*).

The measured fork velocities are shown in Fig. 4B and support the fork-number model: MCH1 replicates the sequences after *crtS* at roughly 1.6 times the fork velocity of the wild-type cells, consistent with the fork-number model. Alternatively, we can consider the same quantitation of fork velocity we considered above: The ratio of fork velocities of loci replicated before *crtS* to those replicated afterwards:

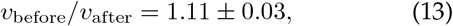

therefore only a very small slowdown is observed after *crtS* is replicated in MCH1, even though exactly the same sequences are replicated, again consistent with the fork-number model.

### The fork velocity oscillates in *E. coli*

Although experiments in *V. cholerae* clearly support the fork-number model, there is significant variability that cannot be explained by this model alone. Are time-dependent variations in fork velocity also observed in organisms that replicate a single chromosome? To answer this question, we worked in the gram-negative model bacterium *Escherichia coli*, which harbors a single 4.6 Mb chromosome. A large collection of marker-frequency datasets have already been generated for both rapid and slow growth conditions by the Rudolph lab [34]. As with the *B. subtilis* marker-frequency datasets, we selected those that had the lowest statistical and systematic noise. (See the Supplemental Material Sec. III C 2.)

The fork velocities are shown in Fig. 5. As before, statistically significant variation is observed in the fork velocity as a function of position. (See Tab. I and Supplemental Material Sec. IV C.) As discussed above in the context of *V. cholerae*, we had initially hypothesized that this variation might be a consequence of rDNA position or some other locus-dependent mechanism; however, there are three arguments against this hypothesis: (i) The slow-velocity regions are not coincident with rDNA locations (Fig. 5A) or relative GC content (Supplement Material Sec. VI A). (ii) Consistent with the time-dependent model, 84% (and 59%) of the observed variation in the fork velocity is symmetric for fast (and slow) growth. (iii) We would expect that a locus-dependent model would predict slow regions that are consistent between fast and slow growth, which is not observed. (See the purple arrows in Fig. 5A.) We therefore conclude that the dominant mechanism for determining the fork velocity is a time-dependent mechanism, consistent with our observations for *V. cholerae*.

**FIG. 5.**
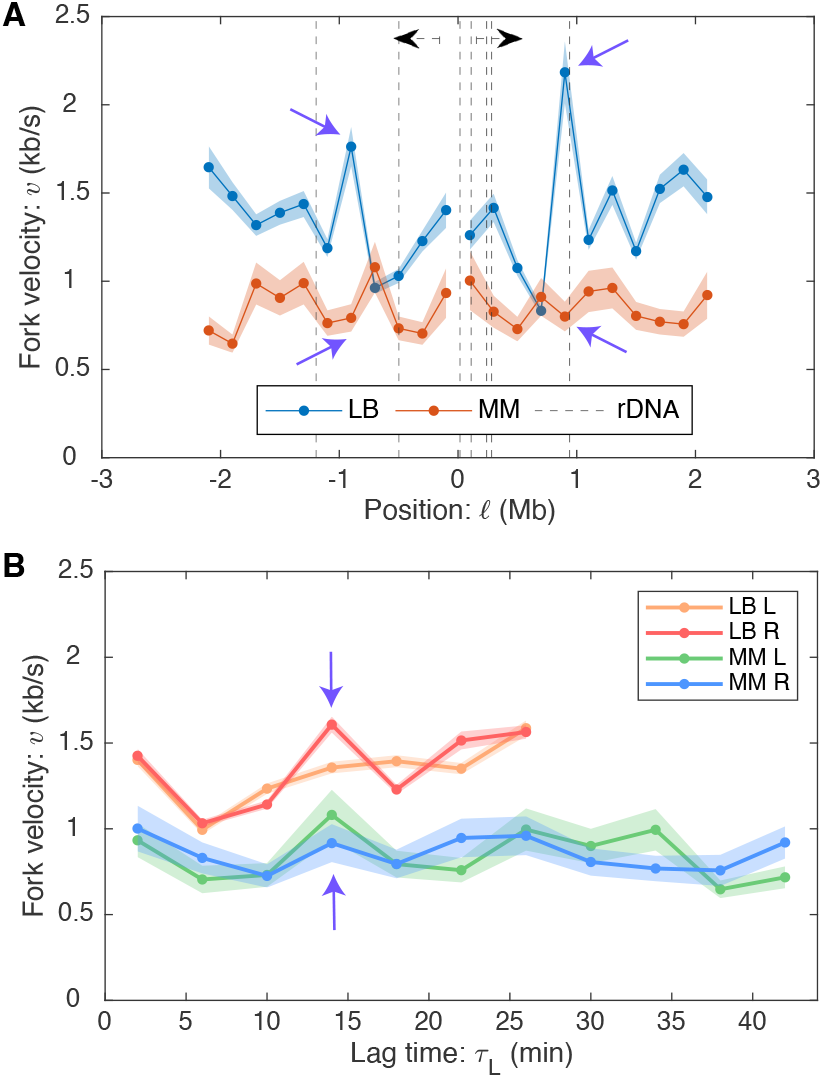
Panel A: Velocity oscillations with respect to position in *E. coli*. We compare fork velocities as a function of genomic position (with respect to *oriC*) under rapid (LB) and slow (minimal media–MM) growth conditions. Motivated by conflict-induced pauses, we have annotated the rDNA positions; however, slow velocities are not consistently coincident with rDNA loci. Regions with high fork velocities are not consistent between rapid and slow growth. *E*.*g*. see the purple arrows. **Panel B: Velocity oscillations with respect to lag time in *E. coli***. The velocity profiles have significant bilateral symmetry: the right and left arm velocities oscillate up and down together. Furthermore, not only are the oscillations consistent between left and right arms, they are also consistent between rapid (LB) and slow growth (minimal media–MM). *E*.*g*. see the purple arrows.

The lag-time analysis is particularly informative in the context of the *E. coli* data (Fig. 5B): Although there is no alignment in the velocity with respect to locus position, there is clear alignment of the fork velocity variation with respect to lag time, not only between the left and right arms of the chromosome, but between slow and fast growth conditions. The observed oscillations appear roughly sinusoidal in character.

### Fork velocity oscillations are observed in three organisms

Temporal oscillations in the fork velocity are an unexpected phenomenon. Are these features a systematic error with a single dataset? First we note that these oscillations are present in two *E. coli* growth conditions (LB and minimal). This phenomenon would be on sounder footing if similar oscillations are observed in two evolutionarily distant species: the gram-negative *V. cholerae* and gram-positive *B. subtilis*. If this phenomenon is observed, to what extent are the oscillations of similar character (*e*.*g*. phase, amplitude, and period)? We compared the lag-time-dependent fork velocity for all three species. In *B. subtilis*, we have already discussed a rDNA-induced pausing on the right arm, which could complicate the interpretation of the data. We therefore consider the fork velocity on just the left arm. For *E. coli* and *V. cholerae*, we compute the average velocity as a function of lag time between the two arms. Since the different organisms and growth conditions have different mean fork velocities, we compare the fork velocity relative to the overall mean. The results are shown in Fig. 6.

**FIG. 6.**
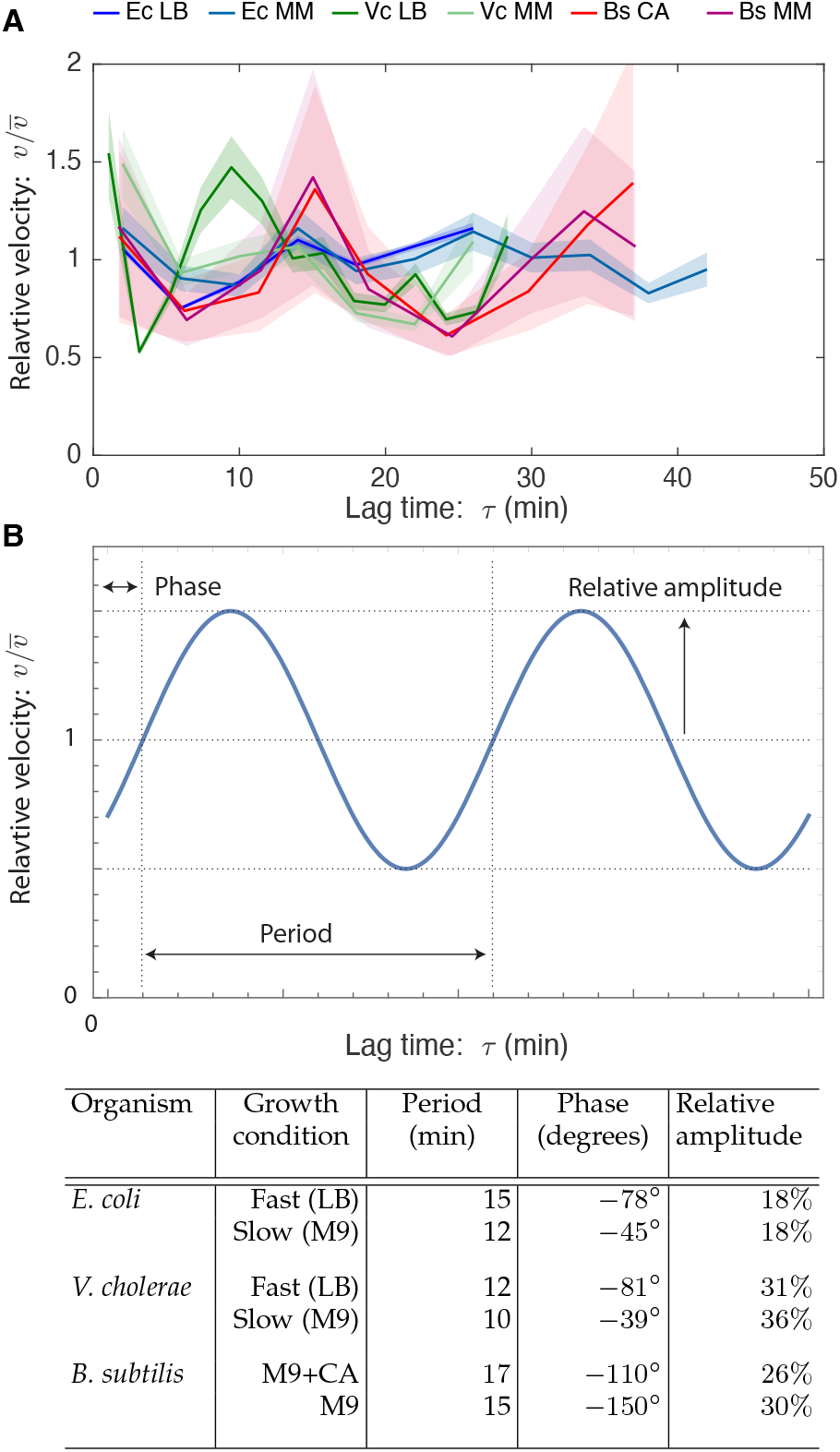
Panel A: Temporal velocity oscillations are observed in three bacterial species: *E. coli* (Ec), *B. subtilis* (Bs), and *V. cholerae* (Vc). The fork velocity starts high before decaying rapidly and then recovering. **Panel B: Oscillation characteristics**. The oscillatory characteristics are broadly consistent both between conditions and species.

All three organisms show oscillations with the same qualitative features: Each fork velocity has roughly the same phase: The velocity begins high, before decaying. The relative amplitudes, roughly 0.5 peak-to-peak, are all comparable with the largest-amplitude oscillations observed in *V. cholerae* and the smallest in *E. coli*. When the relative velocities are compared, it is striking how much consistency there is between growth conditions in *E. coli* and *B. subtilis*. Finally, the period of oscillation is comparable but distinct in all three organisms, ranging from 10 to 15 minutes. The oscillation characteristics are summarized in a table in Fig. 6.

## III. DISCUSSION

The focus of this paper is on the development of lag-time analysis, which uses exponential growth as the timer to characterize cellular dynamics. Although the approach is in principle widely applicable to characterize the dynamics of any molecule of complex, we focus on replication dynamics to explore this novel approach. Although the approach has been understood at a conceptual level since the pioneering work of Cooper and Helmstetter [19], our recent exploration of stochastic models and the introduction of the exponential mean have clarified its interpretation [13] and, from an experimental perspective, the introduction of next-generation sequencing greatly expanded the potential of the approach for characterizing the dynamics of nucleic acids and in particular replication dynamics, where it has the potential to make precise measurements of replication timing. In multiple applications, we have used this approach to quantitatively measure time durations as short as seconds, a time resolution that is challenging, if not impossible, to achieve using other methods.

To appreciate the power of our approach, it is useful to compare our results to recent results of Nieduszynski and coworkers who have used experimental methods, cell synchronization (sync-seq), to directly resolve replication dynamics [35, 36]. In this case, the time resolution is limited by a combination of the precision of cell synchronization, which is an imperfect tool [37], and the frequency of fraction collection (every 5 minutes). Although it would be interesting to compare the relative precision of our approach to this competing method, the authors do not report fork velocities, pause durations, or provide an error analysis of their reported replication times, questioning to what extent the approach is truly quantitative. Since the fractions are collected on five-minute intervals, this time resolution is the floor of the direct time resolution achieved by this approach. In contrast, we report on a range of pause durations that are shorter than 5 minutes.

A significant experimental shortcoming of the sync-seq approach is the necessity of cell synchronization. In most systems, synchronization requires cell-cycle arrest, which introduces a significant potential for artifactual results [14]; whereas lag-time analysis probes dynamics in steady-state growth. Our own preliminary analysis suggests that the timing of initiation at a subset of loci in *Saccharomyces cerevisiae* is changed by the cell synchronization procedure relative to steady-state growth [38].

In addition to these quantitative and high-time-resolution applications, we have also demonstrated the approach in a more conceptual context: using lag-time analysis to argue that the observed oscillations in fork velocity were temporal rather than locus dependent. Together, these examples demonstrate both conceptual and concrete advantages to lag-time analysis over the traditional marker-frequency analysis.

### The significance of the fork velocity

Previous marker-frequency analyses have often reported a log-slope (*e*.*g*. [32, 39]), which is closely related to the fork velocity. What new insights does the measurement of the fork velocity offer over this closely related approach? The fork velocity approach has two important advantages: (i) The first advantage is a conceptual one. The underlying quantity of interest is velocity (or rate per base pair). This is the quantity that is measured *in vitro* and is relevant in a mechanistic model. In contrast, the log-slope is an emergent quantity that is only relevant in the context of exponential growth. (ii) The second advantage is concrete: Although log-slope measurements allow ratiometric comparisons between fork velocity at different loci in the same dataset, *they cannot be used to make comparisons across datasets*. Any comparison of the log-slope between cells with different growth rate (*e*.*g*. due to changes in growth conditions, mutations, species, *etc*.) are meaningless. For instance, the log-slopes of the wild-type and MCH1 *V. cholerae* strains are very different even though the changes in the fork velocity are small. Our wide-ranging comparisons between growth conditions, mutants, and organisms demonstrate the power of reporting fork velocity over the log-slope.

### Applications to eukaryotic cells

Although our focus has been on replication in bacterial cells, an important question is to what extent our approach could be adapted to eukaryotic cells. First, we emphasize that the lag-time analysis is directly applicable without modification to the eukaryotic context. As such, the timing of the replication of loci can be analyzed; however, since the S phase is typically a smaller fraction of the cell cycle and the genomes of eukaryotic cells are larger, deeper sequencing will be required to achieve the same resolution we demonstrate in the context of bacterial cells. One significant potential refinement to this approach is the use of cell sorting (sort-seq) to enrich for replicating cells which can greatly increase the signal-to-noise ratio [35, 36]; however, this approach appears to lead to significant flattening near early-firing origins, as we have observed in other contexts (Supplemental Material Sec. III C 2), and therefore increasing sequencing depth is probably the most promising approach for eukaryotic systems when quantitative characterization is a priority. (See Methods Eqs. 22 and 23 for an estimate of resolution.)

Although lag-time analysis can easily be extended to the eukaryotic context, the measurement of the fork velocity will require some care. A critical assumption in our analysis is that replication forks move unidirectionally at any particular locus, *i*.*e*. it can be either rightward or leftward moving but not both. (See Supplemental Material Sec. III C 9.) Fork traps prevent this bidirectionality in many bacterial cells, but this does not apply to eukaryotic cells. However, if regions of the chromosome can be found where fork movement is unidirectional, *e*.*g*. sufficiently close to early-firing origins, fork velocity measurements could be made in eukaryotic cells. For instance, these conditions appear to be met for a significant fraction of the *Saccharomyces cerevisiae* genome [36]. With significant increases in sequencing depth, we expect analogous replication phenomenology, including pausing and locus- and time-dependent fork velocities, will be observed in eukaryotic systems using lag-time analysis.

### Importance of a model-independent approach

As we prepared this manuscript, we became aware of a competing group which also uses marker-frequency analysis to test a specific hypothesis: the fork velocity is oscillatory in *E. coli* [15], consistent with our own observations. Although our reports share some conclusions, this competing approach requires detailed models for the cell cycle and the fork velocity, along with explicit stochastic simulations. We demonstrate an approach to measure fork velocities *independent of model assumptions* or *detailed hypotheses for the fork velocity*, without the need for numerical simulation and complete with the ability to perform an explicit and tractable error analysis. We therefore expect our analysis will be both widely applicable as well as accessible to other investigators without specialized tools or modeling expertise.

### Systematic error in datasets

It may seem perplexing that we have not pooled many existing datasets from multiple independent experiments. This would naïvely increase the statistical resolution and sensitivity from an analytical perspective. However, it is important to emphasize that not all marker-frequency experiments are of equal quality and that many datasets we have analyzed have clear signatures of systematic error. (See the Supplemental Material Sec. III C 2.) In our analysis, we have prioritized the selection of artifact-free datasets over the indiscriminate pooling of data. We emphasize that to date, datasets have not been generated with quantitative replication dynamics analysis as a goal and we are confident that experimental protocols can be optimized to improve the data. Ref. [40] describes a promising approach, including harvesting populations earlier in exponential phase. We too are developing new protocols to increase data quality.

### Multiple factors determine replisome dynamics

Our measurements of the replication velocity reveal that there are multiple important determinants that results in complex velocity profiles.

### dNTP pools regulate the fork velocity

Previous work had already demonstrated that increases (or decreases) in dNTP pool levels lead to concomitant decreases (or increases) in the C period duration, consistent with a dNTP-limited model of the replication velocity [41–44]. Our data is broadly consistent with these previous results, but in a subcellular context: (i) The fork-number model, in which fork velocities decrease as the number of active forks increase, is clearly consistent with a mechanism in which the nucleotide pool levels, although highly-regulated [45], cannot completely compensate for the increased incorporation rate associated with multiple forks. (ii) The observation of the fork velocity oscillations is also consistent with an analogous failure of the regulatory response to compensate, this time temporally. The initial fall in the fork velocity is consistent with a model in which dNTP levels initially fall as replication initiates and nucleotides begin to be incorporated into the genome. Reduction in the dNTP levels causes a regulatory response to increase dNTP synthesis by ribonucleotide reductase [45], but the finite response time of the network could lead to dynamic overshoot in the regulatory feedback, leading to oscillations [46]. Ref. [15] has also argued that this oscillating-dNTP-level model would lead to time-dependent oscillations in the mutation rate which are consistent with the origin-mirror-symmetric distribution of the mutation observed in *E. coli*. However, this interesting phenomenon and this hypothesized mechanism will require further investigation.

### Retrograde fork motion leads to slow replication velocities

Retrograde fork motion, where the fork moves in the opposite direction from wild-type cells, lead to the largest changes in fork velocity observed. To what extent is the observed slowdown a consequence of a few long-duration pauses versus a region-wide slowdown? In regions which exclude the rDNA, the effect appears well distributed. However, it is important to note that the genomic resolution of lag-time analysis is still much too low to resolve individual transcriptional units. We anticipate that with increased sequencing depth as well as improvements in sample preparation, this approach could detect genomic structure in the fork velocity at the resolution of individual transcriptional units. Although we did analyze a number of mutants with retrograde fork movement in *V. cholerae* and *E. coli* (analysis not shown), the competing effect of increased fork number as well as the genomic instability of these strains made these experiments difficult to interpret quantitatively, since fork number and direction were both affected in these strains [32, 47]. We concluded qualitatively that retrograde replication direction appears not to play as large a role in these gram-negative bacteria as it does in gram-positive *B. subtilits*, consistent with previous evidence [31, 32, 48–50]. However, we expect lag-time analysis could be used to characterize even small effects of the retrograde fork orientation in more-carefully engineered strains, analogous to those that we analyzed in the context of *B. subtilis* [31, 32].

Previous reports [31, 32], including our own [12, 51– 54], had reported long-duration replication-conflict induced pauses, especially in mutant strains where the orientation of rDNA [31] or other highly transcribed genes [51] are inverted to give rise to a head-on conflict between replication and transcription. The contribution of lag-time analysis in this context is multifold: First, we provide a quantitative number in the context of the very-short-duration pauses for co-directional transcription in wild-type cells. This analysis supports a longstanding hypothesis that the right arm of the *B. subtilis* chromosome is shorter than the left arm to compensate for pausing at the rDNA loci that arm predominately located on this arm.

We also report quantitative measurements for the longer pauses that results from head-on conflicts in mutants where highly transcribed genes are inverted. Our analysis gives us the ability to quantitatively differentiate the contributions of fork pausing and arrest in the analysis of the marker frequency, which was previously impossible. Our measurement of a timescale of minutes is consistent with our previous *in vivo* single-molecule measurements in which we report transcription-dependent disassembly of the core replisome [12]. Could the observed fork-velocity oscillations be misinterpreted as pauses? The observed lag-time offset between the two arms (*e*.*g*. Fig. 3B) is not predicted by fork-velocity oscillations.

### Conclusion

In this paper, we introduce a novel method for quantitively characterizing cellular dynamics by lag-time analysis. Although more broadly applicable, we focus our analysis on the characterization of replication dynamics using next-generation sequencing to quantitate DNA locus copy number genome-wide. The approach has the ability to make precise, even at the resolution of seconds, measurements of time differences and pause durations, as well as the ability to quantitatively measure fork velocities *in vivo* in physiological units of kb/s, at genomic resolutions of roughly 100 kb, for the first time. Importantly, unlike marker-frequency analysis, our approach allows direct quantitative comparisons to be made between growth conditions, mutant strains, and even different organisms. The resulting measurements of replication dynamics reveal complex phenomenology, including temporal oscillations in the fork velocity as well as evidence for multiple mechanisms that determine the fork velocity. The lag-time analysis has great potential for application beyond bacterial systems as well as the potential to significantly increase in resolution and sensitivity as sequencing depth and sample preparation improve.

## IV. METHODS

### Introduction to marker-frequency analysis

Our focus will be on marker-frequency analysis, which measures the total number of a genetic locus in an asynchronous population. The model was generalized to predict the marker frequency *N* (𝓁) of a locus a genomic distance 𝓁 away from the origin [20–22]:

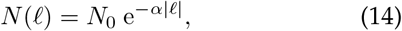

where *N*_0_ is the number of origins, which grows exponentially in time with the rate of mass doubling of the culture, *k*_*G*_. Since the origin is replicated first, the number of origins is always largest compared to the numbers of other loci. Quantitatively, the copy number is predicted to decay exponentially with log-slope:

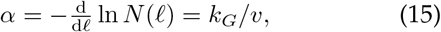

where *k*_*G*_ is the population growth rate and *v* is the fork velocity, typically expressed in units of kilobases per second. To derive this result, two critical assumptions were made: (i) the timing of the cell cycle is deterministic and (ii) the fork velocity is constant [19, 20].

Initially, our naïve expectation was that the interplay between the significant stochasticity of the cell-cycle timing with the asynchronicity of the culture would prevent marker-frequency analysis from being used as a quantitative tool for characterizing cell-cycle dynamics. For instance, significant stochasticity is observed in the duration of the B period [55] (*i*.*e*. the duration of time between cell birth and the initiation of replication). Does this stochasticity lead to a failure of the log-slope relation (Eq. 15)?

### Stochastic simulations support the universality of the log-slope relation

To explore the role of stochasticity and a locus-dependent fork velocity in shaping the marker frequency, we simulated the cell cycle using a stochastic simulation. Our aim was not to perform a simulation whose mechanistic details were correct, but rather to study how strong violations of the Cooper-Helmstetter assumptions, in particular how stochasticity, as strong or stronger than that observed, influenced the observed marker frequency and the log-slope relation (Eq. 15). In short, we used a Gillespie simulation [56] where the B period duration and the lifetime of replisome nucleotide incorporation steps are exponentially distributed, and we added regions of the genome where the incorporation rate was fast as well as a single slow step on one arm. See Fig. 7A and the Supplemental Material Sec. V A for a detailed description of the model, as well as movies of the marker frequency approaching steady-state growth, starting from a single-cell progenitor.

**FIG. 7.**
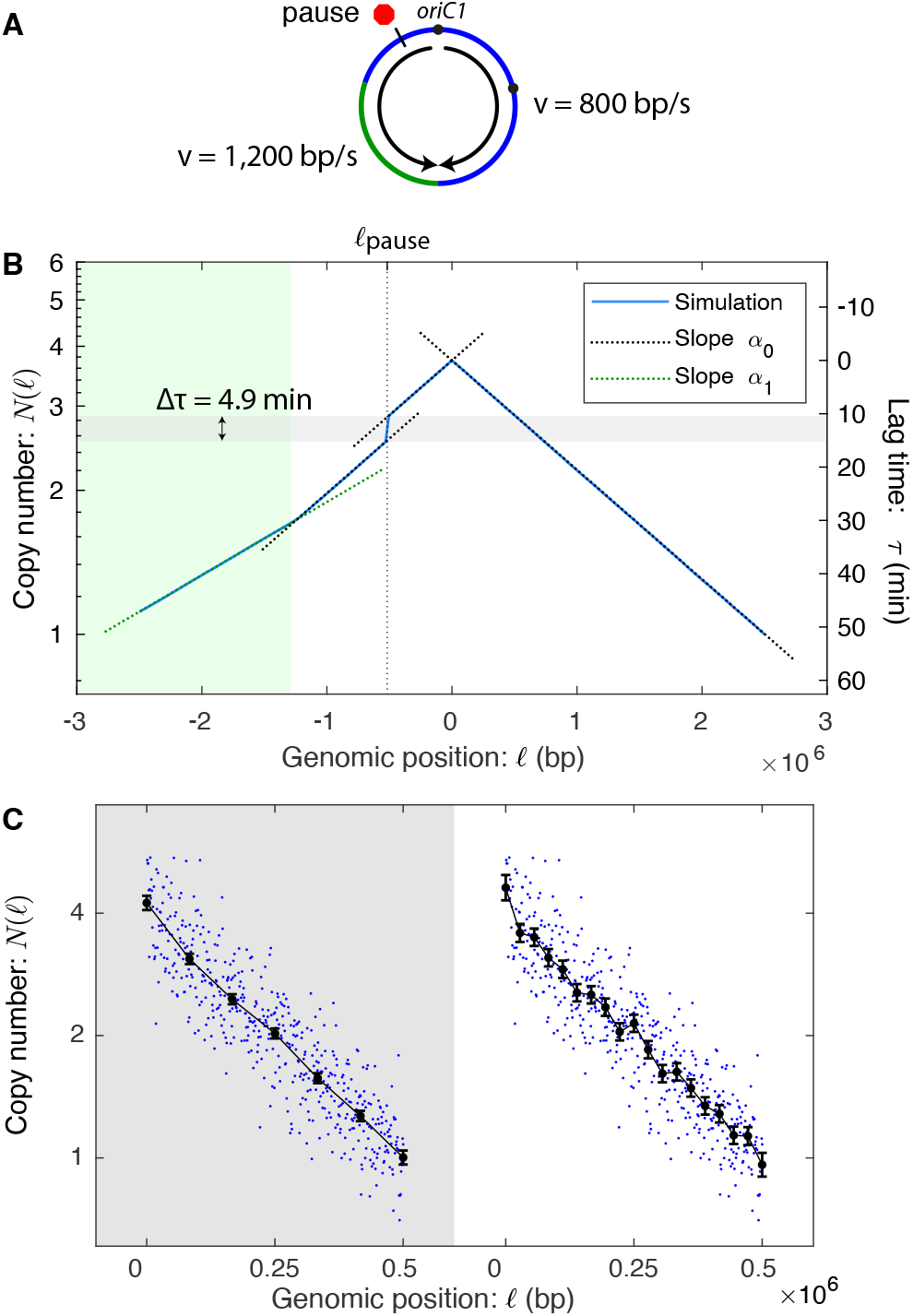
Panel A: A schematic of the simulated chromosome. Replication initiates at the origin, pauses at a locus (red octagon) on the left arm and the velocity is increased on the lower left arm (green). **Panel B: Simulated marker frequency obeys the log-slope law**. The stochastic simulation generates a marker-frequency curve (blue). The model is stochastic in the timing of replication initiation as well as the fork dynamics and it includes two regions (blue and green) with different fork velocities as well as a pause with a stochastic lifetime. (See the *Terminus 4* model in the Supplemental Material Sec. V A.) In spite of the stochasticity, it obeys the log-slope law locally (Eq. 16). Furthermore, the inferred lag-time pause (4.9 min) is predicted by the exponential mean (Eq. 2). **Panel C: Tradeoff between genomic resolution and velocity precision**. As the spacing between knots decreases, increasing the genomic resolution, the error in the velocity measurement increases.

To our initial surprise, the stochasticity of the model had no effect on the predicted log-slope of the locus copy number. (See Fig. 7B.) In spite of the stochastic duration of the B period and the locus-dependence, the marker frequency still decays exponentially with the same decay length *locally, i*.*e*.:

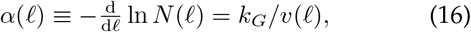

where *k*_*G*_ was the empirically determined growth rate in the simulation and *v*(𝓁) was the local fork velocity at the locus with position 𝓁.

We hypothesized that this result might be a special case of choosing an exponential lifetime distribution, since this is consistent with a stochastic realization of chemical kinetics. To test this hypothesis, we simulated several different distributions, including a uniform distribution, for the duration of the B period and the stepping lifetime for the replisome (as well as simulating multifork replication). In each case, the local log-slope relation (Eq. 16) held, even as the growth rate and fork velocities changed with the changes in the underlying simulated growth dynamics. We therefore hypothesized that Eq. 16 was a universal law of cell-cycle dynamics and independent of Cooper and Helmstetter’s original assumptions.

### The exponential-mean duration

Motivated by this empirical evidence, we exactly computed the population demography in a class of stochastically-timed cell models. We described these results elsewhere [13]. In short, we showed that there is an exact correspondence between these stochastically-timed models and deterministically-timed models in exponential growth. The relationship between the corresponding deterministic lifetime *τ*_*i*_ of a state *i* and the underlying distribution *p*_*i*_ in the stochastic model is the exponential mean (Eq. 2) [13]. The exponential mean biases the mean towards short times, the growth rate *k*_*G*_ determines the strength of this bias, and the biological mechanism for this bias is due to the enrichment of young cells relative to old cells in an exponentially growing culture [13].

To understand the consequences of this result, we consider two special cases of this exponential mean. For processes with lifetimes short compared to the doubling time, Eq. 2 can be Taylor expanded to show that the exponential mean is:

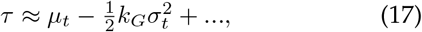

the regular arithmetic mean *μ*_*t*_ with a leading-order correction proportional to the product of the growth rate and variance 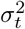. In the context of single-nucleotide incorporation, this correction is on order one-part-in-amillion and therefore can be ignored. As a consequence, Eq. 16, corresponding to the transitions between states with short-lifetimes, is unaffected by the stochasticity, exactly as we observed in our simulations.

Another important case to consider is the strong disorder limit, in which a small fraction of the population *ϵ* stochastically arrests, *i*.*e*. with lifetime ∞, while the other individuals have exponential-mean lifetime *τ*_0_. Using the definition in Eq. 2, it is straightforward to show that the deterministic lifetime is:

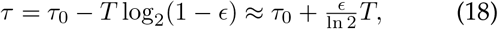

where *T* is the population doubling time and the second equality is an approximation for small *ϵ*. The exponential mean duration is extended by the arrest, but remains finite. Therefore, an arrest of a subpopulation is indistinguishable from a longer duration pause in an exponentially proliferating population. (See Ref. [13].)

### Marker-frequency demography

For a stochastic model with locus-dependent fork velocity, we showed that Eqs. 14 and 15 generalize to

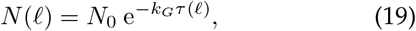

where we will call *τ* (𝓁) the lag time of a locus at position 𝓁, which is equal to the sum of the differential lag times for each sequential step:

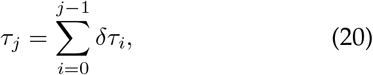

where *δτ*_*i*_ is the differential lag time for state *i* or the exponential mean of the state lifetime [13]. In the continuum limit, it is more convenient to represent this sum as an integral:

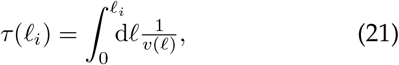

where the fork velocity is defined: *v*(𝓁_*i*_) ≡ 1 bp*/δτ*_*i*_. To demonstrate that the generalized stochastic model predicts the log-slope relation (Eq. 16), the log-slope can be derived by substituting Eq. 21 into Eq. 19, as was observed in the stochastic simulations, demonstrating the universality of Eq. 4. We note that Wang and coworkers had previously derived an equivalent expression using the deterministic framework of the Cooper-Helmstetter model in the Material and Methods Section of Ref. [31].

### Stochasticity has a minimal effect on the marker frequency

We initially had hypothesized that stochasticity should affect the marker frequency. As explained above, it is the rapidity of base incorporation that explains why stochasticity is dispensable in this context. The same argument does not apply to the B period which is comparable to the duration of the cell cycle. However, for the marker frequency, it is lag-time differences between the replication times of loci that is determinative, and therefore the lag time of the B period cancels from these lagtime differences. Although it is mostly irrelevant for understanding wild-type cell dynamics, stochasticity and an arrested subpopulation will play an important role in one phenomenon we analyze: replication-conflict induced pauses.

### Time resolution

Due to the large number of reads achievable in next generation sequencing, the time resolution will be high in carefully designed analyses. The number of reads is subject to counting or Poisson noise. It is therefore straightforward to estimate the experimental uncertainty in the lag time due to finite read number:

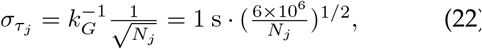

where we have used a read number inspired by the replication-conflict pausing example. This estimate suggests that under standard conditions, time measurements with an uncertainty of seconds are possible using this approach.

### Fork-velocity resolution

To compute the slope in Eq. 4, the log-marker-frequency is fit to a piecewise linear function with equal spacing between knots. See Fig. 7B. There is an important tradeoff between genomic resolution (*i*.*e*. the genomic distance between knots) and fork velocity precision (*i*.*e*. the uncertainty in velocity measurement): Increasing the genomic distance between knots reduces the genomic resolution but also reduces the uncertainty in the velocity measurement. We therefore consider a series of models with increasing genomic resolution and use the Akaike Information Criterion (AIC) to select the optimal model [28, 29]. See Supplemental Material Sec. III C 10. This approach balances the desire to resolve features by increasing the genomic resolution with the loss of velocity precision.

Given a knot spacing, it is straightforward to estimate the relative error:

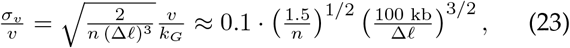

where *n* is the read depth in reads per base and Δ𝓁 is the spacing between knots in basepairs. Therefore, for a canonical next-generation-sequencing experiment, we can expect to achieve roughly 10% error in the fork velocity for 100 kb genomic resolution. Note that in our error analysis, we have included only the error from cell number *N*, not the error from the uncertainty in the cellcycle duration, which covaries between loci in a particular experiment.

## Acknowledgments

The authors would like to thank B. Traxler, A. Nourmohammad, J. Mougous, and J. Mittler for many useful conversations. We would like to thank P. Levin, J. Wang and L. Simmons for advice on our manuscript. We thank S. B. Peterson and A. Schaefer for help with *V. cholerae*. We would like to thank J. Wang, C. Possoz, F.-X. Barre, C. Rudolph for detailed conversations about their data. This work was supported by NIH grant R01-GM128191.

## Data availability statement

The sequencing datasets generated during the current study are currently available on reasonable request, but will be uploaded to a public repository prior to publication. Relevant sequencing data from other studies can be found at the accession numbers given in Sec. II of the Supplemental Material. Digitized data are available on reasonable request.

## Code availability statement

All MATLAB scripts written for this study are available on reasonable request.

