## Supplemental Materials for "The high-resolution *in vivo* measurement of replication fork velocity and pausing by lag-time analysis"

#### CONTENTS

I. Strains used in this study S2

II. Datasets used in this study S3

III. Experimental methods S3
A. Growth media and determination of growth phase S3
B. Generation of marker frequency data for this study S4
C. Marker frequency analysis S4
1. Sequence alignment S4
2. Data selection S4
3. To divide or not to divide by stationary phase data S5
4. Filtering the data S5
5. Nearest-neighbor variance estimator S5
6. Fitting slopes to the log of the copy number S6
7. Estimating errors for the control points S6
8. Transformation matrices for obtaining the fork velocity S7
9. Behavior near the terminus S7
10. Selection of the number of fitted knots S7

IV. Theoretical methods S8
A. Bilateral symmetry analysis S8
B. Estimation of average fork number per cell cycle S8
C. Statistically significant deviations of local fork velocity from the global mean S10

V. Supplemental theoretical results S10
A. Stochastic simulations S10
1. Stochastic simulations method S10
2. Stochastic simulations match the predictions of the log-slope S11
3. Simulation models S11
4. Simulation results S12
5. Re-scaling simulation units to compare to measured data S12
6. Movies of marker frequency dynamics approaching steady state growth S12

VI. Supplemental experimental results S13
A. Comparison with GC content S13
B. *V. cholerae* WT on LB S14

*

†; http://mtshasta.phys.washington.edu/

C. *V. cholerae* WT on M9 fructose S15
D. *V. cholerae* MCH1 on LB S16
E. *V. cholerae* MCH1 on M9 fructose
F. *V. cholerae oriR4* on M9 fructose S17
G. *B. subtilis oriC*-257° on S7 fumarate S18
H. *B. subtilis oriC*-94° on S7 fumarate S18
I. *B. subtilis oriN* on S7 fumarate S19
1. Two-slopes model S19
2. Pause model S19
J. *B. subtilis oriN*-257° on S7 fumarate S20
K. *B. subtilis rrnIHG* inversion on MOPS glucose w/ Casamino Acids S20
L. *B. subtilis rrnIHG* inversion on MOPS glucose – Minimal S21
M. *E. coli* WT on LB S22
N. *E. coli* WT on M9 glucose S22

References S23

#### I. STRAINS USED IN THIS STUDY

| Short name: | Strain name: | Description: | Source: |
|---|---|---|---|
| *V. cholerae* WT | O1 biovar El Tor N16961 | *ChapR* Δ*lacZ* gm$^R$ | [1] |
| *V. cholerae* MCH1 | MCH1 | Integration of N16961 Chr2, without *oriC2* and partition machinery region, in place of the *dif1* site in N16961 Chr1. | [1] |
| *V. cholerae oriR4* | EGV111 | N16961 *ChapR* Δ*lacZ oriC1* @ R4 (1,898Mb) gm$^R$ | [1] |
| *B. subtilis* WT | JH642 | *trpC2 pheA1* | [2] |
| *B. subtilis* 257°::*oriC* | MMB703 | *trpC2 pheA1 argG*(257°)::(*oriC*/*dnaAN kan*) Δ(*oriC-L*)::*spc* | [2] |
| *B. subtilis* 94°::*oriC* | JDW258 | *trpC2 pheA1 aprE*(94°)::(*oriC*/*dnaAN kan*) Δ(*oriC-L*)::*spc dnaB134ts-zhb83*::Tn*917*(*cat*) | [2] |
| *B. subtilis oriN* | MMB208 | *pheA1* (*ypjG-hepT*)*122 spoIIIJ*(359°)::(*oriN kan tet*) Δ*oriC-S* | [2] |
| *B. subtilis* 257°::*oriN* | MMB700 | *pheA1* (*ypjG-hepT*)*122 argG*(257°)::(*oriN kan*) Δ*oriC-S* | [2] |
| *B. subtilis* YB886 | YB886 | *trpC2 metB5 sigB amyE spβ-ICEBs° xin-* | [3] |
| *B. subtilis rrnIHG*(pre-inv) | JDW858 | YB886 *kbaA'*::*neo' cat*::*'ybaN rrnG-5S'*::*erm 'neo*::*'ybaR* | [3] |
| *B. subtilis rrnIHG*(inv) | JDW860 | YB886 *kbaA'*::*neo erm* inv(*ybaN*::*rrnG-5S*) *cat*::*'ybaR* | [3] |
| *E. coli* WT | K-12 MG1655 | F- lambda- *ilvG- rfb*-50 *rph*-1 | [4] |

#### II. DATASETS USED IN THIS STUDY

All datasets were obtained from cells grown at 37 °C. Next-generation-sequencing FASTQ files and marker frequencies were either generated for this study, obtained from Wang et al. [2, 3], or obtained from the European Nucleotide Archive (ENA). The corresponding project accessions are included if applicable. In the following table, Exp is shorthand for exponential phase growth and Stat is shorthand for stationary phase. Growth media are described in more detail in Supplemental Material Sec. III A.

| Short name: | Growth media: | Doubling time: | Source: | Project accession: | Run accession: |
|---|---|---|---|---|---|
| *V. cholerae* WT | LB | 22 ± 1 min | This study | TBA | Exp: 8217-AJ-013<br>Stat: 8217-AJ-001 |
| | M9 fructose | 50 ± 4 min | From study [1] | ENA: PRJEB28538 | Exp: ERX2796386<br>Stat: ERX2796387 |
| *V. cholerae* MCH1 | LB | 90 ± 6 min | This study | TBA | Exp: 8217-AJ-016<br>Stat: 8217-AJ-004 |
| | M9 fructose | 50 ± 4 min | From study [1] | ENA: PRJEB28538 | Exp: ERX2796384<br>Stat: ERX2796385 |
| *V. cholerae oriR4* | M9 fructose | 55 ± 5 min | From study [1] | ENA: PRJEB28538 | Exp: ERX2796379<br>Stat: ERX2796380 |
| *B. subtilis* WT | S7 fumarate | 61.0 ± 1.2 min | From study [2] | Digitized | Digitized |
| *B. subtilis* 257°::*oriC* | S7 fumarate | Not reported | From study [2] | Digitized | Digitized |
| *B. subtilis* 94°::*oriC* | S7 fumarate | Not reported | From study [2] | Digitized | Digitized |
| *B. subtilis oriN* | S7 fumarate | 64.3 ± 2.9 min | From study [2] | Digitized | Digitized |
| *B. subtilis* 257°::*oriN* | S7 fumarate | Not reported | From study [2] | Digitized | Digitized |
| *B. subtilis rrnIHG*(pre-inv) | LB | 20 ± 1 min | From study [3] | Digitized | Digitized |
| | MOPS glucose CA | 28 ± 1 min | From study [3] | Digitized | Digitized |
| | MOPS glucose MM | 42 ± 1 min | From study [3] | Digitized | Digitized |
| *B. subtilis rrnIHG*(inv) | LB | > 160 min | From study [3] | Digitized | Digitized |
| | MOPS glucose CA | 44 ± 5 min | From study [3] | Digitized | Digitized |
| | MOPS glucose MM | 44 ± 1 min | From study [3] | Digitized | Digitized |
| *E. coli* WT | LB | 19.3 ± 1.7 min | From study [4] | ENA: PRJEB25595 | Exp: ERS2298483<br>Stat: ERS2298484 |
| | M9 glucose | 68.8 ± 6.2 min | From study [4] | ENA: PRJEB25595 | Exp: ERS2298504<br>Stat: ERS2298505 |

#### III. EXPERIMENTAL METHODS

##### A. Growth media and determination of growth phase

As we have used data from multiple sources, the minimal media and the determination of population growth phase are not consistent across all studies. All studies use the standard recipe for Luria-Bertani (LB) rich medium (1% tryptone, 0.5% yeast extract, and 1% NaCl in H2O), with the exception of Midgley-Smith et al. (2018), where 0.05% NaCl is used instead. All studies using M9 minimal media have the same base recipe (1X M9 salts, 2 mM MgSO4, and 0.1 mM CaCl2), with different supplements and carbon sources. In our study and in Galli et al. (2019) [1], M9 was supplemented with 10 $\mu$g/mL thiamine HCl for the *V. cholerae* strains. The carbon source for M9 is 0.4% glucose for our study, 0.4% fructose for Galli et al. [1], and 0.2% glucose for Midgley-Smith et al. [4]. In Wang et al. (2007) [2], the relevant datasets exclusively use S7 minimal medium (50 mM MOPS, 10 mM (NH4)2SO4, 5 mM potassium phosphate, 2 mM MgCl2, 0.9 mM CaCl2, 50 $\mu$M MnCl2, 5 $\mu$M FeCl3, 10 $\mu$M ZnCl2, and 2 $\mu$M thiamine hydrochloride), supplemented with 1% sodium fumarate as the carbon source, 0.1% glutamate, 40 $\mu$g/mL tryptophan, and 40 $\mu$g/mL phenylalanine. For Srivatsan et al. (2010) [3], the minimal medium consists of 50 mM MOPS with 1% glucose, along with different supplements for the CA (0.5% casamino acids) and MM (40 $\mu$g/mL tryptophan, 40 $\mu$g/mL methionine, 40 $\mu$g/mL phenylalanine, and 100 $\mu$g/mL arginine) growth media.

All populations were grown at 37 °C. In our study, exponential phase cultures were grown to $OD_{600}$ of 0.60-0.80. Stationary phase samples were grown to $OD_{600}$ of 1.50 or greater. In Galli et al. [1], exponential phase cultures were grown to an $OD_{650}$ of 0.05 and 0.2 in M9 and LB, respectively. In Midgley-Smith et al. [4], exponential phase cultures were grown to an $OD_{600}$ of 0.48. In Srivatsan et al. [3], exponential phase cultures were grown to an $OD_{600}$ of 0.2-0.6.

##### B. Generation of marker frequency data for this study

For detailed protocols of marker-frequency generation for each dataset, see the corresponding references in Sec. II. In this study, strains were struck on solid agar and grown overnight at 37 °C. Three individual colonies were selected from each strain and used to inoculate 10 mL of LB media, which was grown overnight at 37 °C w/ 260 rpm shaking. The following morning, these overnight pre-cultures were back-diluted to $OD_{600}$ of 0.05 in 5 mL of either LB or M9 media. The same biological replicate was used to inoculate both growth conditions. Exponential phase cultures were grown to $OD_{600}$ of 0.60-0.80. Stationary phase samples were grown to $OD_{600}$ of 1.50 or greater. To harvest, cultures were centrifuged at 10,000 rpm for 5 mins at 25 °C, and the supernatant was removed. gDNA was prepared immediately following harvest using GeneJet Genomic DNA Purification Kit (Thermo Scientific, K0721). Purified gDNA was quantified using Qubit dsDNA HS Assay Kit (Invitrogen, Q32851) and quality was assayed using a Nanodrop 2000 Spectrophotometer (Thermo Scientific, ND-2000). Whole-genome libraries were prepared from purified gDNA using Twist 96-Plex Library Prep Kit (Twist Bioscience, 104950) with dual adapters. Libraries were subsequently pooled and sequenced on Illumina NovaSeq6000 (S4) to a depth of 6,000,000 reads per sample.

##### C. Marker frequency analysis

###### *1. Sequence alignment*

To align deep sequencing read outputs to the bacterial reference genomes for MFA, we used the read alignment tool Bowtie 2 via the MATLAB function `bowtie2` [5]. We then extracted the position information from the resulting SAM file using an in-lab written MATLAB script and the command-line tool SAMtools [6].

###### *2. Data selection*

Not all marker-frequency experiments are of the same quality and many datasets we have analyzed have clear signatures of systematic error. We use two key criteria to determine the quality of the data. The first is a qualitative measure involving the shape of the marker-frequency profile and the second is a quantitative measure based on resolution analysis.

**Marker profile near origin.** Our analysis uses exponential growth as the stop-watch for resolving dynamics, so it is essential that the population is harvested at the right time for sequencing. We have found that the marker-frequency profile has a signature flattening near the origin of replication if the population is harvested too late in exponential phase, as shown in Fig. 1. The population is entering stationary phase, where nutrient depletion begins to cause slower growth. Entering stationary phase also causes a decrease in the population-wide rate of replication initiation, which leads to the flattening of the profile near the origin. For the rest of the chromosome, replication continues unchanged for the majority of the population. Since exponential growth is crucial to our analysis, we have chosen to only use datasets with cusp-like behavior near the origin(s), as opposed to a rounded concave-down shape.

**Resolution analysis.** In the case where there is a single origin of replication for each chromosome, we expect only a single maximum in the marker frequency profile, corresponding to the origin. In an ideal noiseless situation, any local maxima distinct from the origin would correspond to other points of replication initiation and the fit would predict a retrograde fork velocity. However, due to the stochastic nature of the sequencing data, there can be spurious local maxima if the resolution is chosen to be too high. Fluctuations due to noise can cause the knots in the piecewise-linear fit along each arm of the chromosome to be non-monotonic, as shown in Panel C of Fig. **??** in the main paper. Thus, given two datasets for the same organism and growth condition, data quality was determined based on the best resolution that can be achieved without introducing spurious maxima.

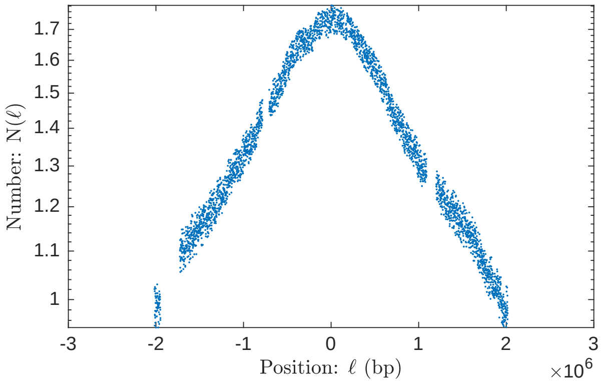


FIG. 1. Marker frequency profile of *V. cholerae* MCH1 in LB rich media. Flattening of the profile near the origin of replication suggests a slowdown in the rate of replication initiation, consistent with the population entering stationary phase.

##### 3. *To divide or not to divide by stationary phase data*

To further remove any sequencing induced bias, we divide the copy number data obtained during exponential growth by the data obtained during stationary phase. This normalization step is done with the 1 kb binned values. Since the division of two noisy uncorrelated data sets typically results in a decreased signal-to-noise ratio (SNR), we used a comparison of the variance divided by the mean before and after division by stationary phase.

We expect the relative variance to decrease by normalization if the reduction in sequencing bias resulting from division has a larger effect than the increase in noise from dividing two noisy data sets. We first divide the noisy stationary phase data by its expected mean value. This ensures that it is of unit order and will not introduce a global scale factor during division, which would affect only the normalized data and not the pre-normalization data. We find that after division, the variance is roughly halved, suggesting that the normalization by stationary phase is successful in reducing sequencing bias without resulting in excessive noise. We thus chose to divide all exponential phase data with the corresponding stationary phase data. For some datasets, the stationary phase results were not provided for some of the ectopic origin mutants. The only difference in these mutants from their original strains is the addition of a short segment of DNA for the ectopic origin, which does not significantly change the locations of sequencing bias. Therefore, we chose to divide the exponential phase data for the ectopic origin strains by the stationary phase data from the original strains.

##### 4. *Filtering the data*

To remove outliers from the marker-frequency data, we used a centered 200 kb median filter, such that at each position $\ell$ along the chromosome, we set the median marker-frequency from $\ell - 100$ kb to $\ell + 100$ kb as a baseline. The data was given periodic boundary conditions such that the centered medians still had 200 kb regions at the boundaries of the data set. We then discarded the 10% of points that most deviated from this centered median baseline, sorted by the absolute value of the difference. We also removed data points from regions known to be associated with mobile elements of low GC content, such as the Super Integron on Chr2 of *V. cholerae*, which is subject to anomalous sequencing bias.

##### 5. *Nearest-neighbor variance estimator*

We used a nearest-neighbor estimate of variance:

$$\hat{\sigma}^2 \approx \frac{1}{N}\sum_{i=1}^{N}\frac{1}{2}(x_{i+1} - x_i)^2, \tag{1}$$

where $x_{N+1} = x_1$, in agreement with periodic boundary conditions. This approach allows us to obtain a variance measurement without assuming the underlying distribution of the data and the corresponding expected value of each bin.

##### 6. *Fitting slopes to the log of the copy number*

After taking the natural logarithm of the copy number, we expect the data to follow a roughly linear trend along each arm of the chromosome. Since our goal is to obtain higher resolution measurements of fork velocity along the chromosome, it is necessary to subdivide each arm into smaller segments with variable slopes. The fit to the data should be continuous, since discontinuities would create copy number ambiguity at segment junctions. Therefore, we use a continuous piecewise linear least squares fit to obtain slope measurements from the data. We call the MATLAB function lsqnonlin as part of an in-lab written fitter function. The fitter obtains vertical control point values at the junctions between segments. Each control point is used in the measurement of the slope both to the left and to the right of it.

##### 7. *Estimating errors for the control points*

To obtain error estimates for the control points of the least squares fit, we use the Jacobian that lsqnonlin returns, which is the Jacobian of the difference between the fit data and the experimental data. If we have $k$ fit parameters $\theta_\alpha$, where $\alpha = 1, ..., k$, and $N$ data points $y_i$, where $i = 1, ..., N$, then the Jacobian takes the form:

$$J_{i\alpha} = \frac{\partial}{\partial \theta^\alpha}(y_i - \mu_i(\theta)), \tag{2}$$

where $\mu_i(\theta)$ is the fitter estimate of $y_i$, based on the model with the set of parameters $\theta$. We use the matrix form of the Fisher information:

$$[I(\theta)]_{\alpha,\beta} = \mathbb{E}\left[\left(\frac{\partial}{\partial \theta^\alpha}\log f(Y;\theta)\right)\left(\frac{\partial}{\partial \theta^\beta}\log f(Y;\theta)\right)\middle|\theta\right], \tag{3}$$

where $f(Y;\theta)$ is the probability density function (PDF) of the random variable $Y$, given parameters $\theta$. In our case, we expect the noise to be Gaussian distributed around the mean, so we have the PDF:

$$f(Y;\theta) = \frac{1}{(\sigma\sqrt{2\pi})^N}\prod_{i=1}^{N}\exp\left(-\frac{(y_i - \mu_i(\theta))^2}{2\sigma^2}\right), \tag{4}$$

where $\sigma$ is the standard deviation. Now we take the derivative of the logarithm:

$$\frac{\partial}{\partial \theta^\alpha}\log f(Y;\theta) = \frac{\partial}{\partial \theta^\alpha}\left(-\frac{\sum_i (y_i - \mu_i(\theta))^2}{2\sigma^2} - N\log\left(\sigma\sqrt{2\pi}\right)\right), \tag{5}$$

$$= -\frac{1}{\sigma^2}\sum_{i=1}^{N}(y_i - \mu_i(\theta))\frac{\partial}{\partial \theta^\alpha}(y_i - \mu_i(\theta)), \tag{6}$$

$$= -\frac{1}{\sigma^2}\sum_{i=1}^{N}(y_i - \mu_i(\theta))J_{i\alpha}. \tag{7}$$

Plugging this into Eq. 3 and noting that the matrix multiplication contracts over the $i$ index, we are left with:

$$I(\theta) = \frac{J^T J}{\sigma^2}. \tag{8}$$

The Cramér-Rao bound states that the inverse of the Fisher information provides a lower bound for the covariance matrix of unbiased estimators. More explicitly, the relation is as follows:

$$\mathrm{cov}_\theta\left(\hat{\theta}\right) \geq I(\theta)^{-1}. \tag{9}$$

We take the equality for our error estimates. The variances of the parameter estimates are given by the diagonal elements of the covariance matrix.

##### 8. *Transformation matrices for obtaining the fork velocity*

To convert the control points of the piecewise linear fit into slopes and fork velocities, we use coordinate transformation matrices, which have the added benefit of also transforming the variance estimates from the Fisher information. The coordinate transformation matrices can either be obtained through direct reasoning or as Jacobians of the final coordinates relative to the initial coordinates. The following examples will be square matrices of dimension 3, acting on sets of 3 parameters. These examples illustrate the procedure of obtaining the transformation matrices and are easily generalizable. To convert from a vector of control point values to a vector where the first element is the leftmost control point value and the other elements are the slopes, we use the following matrix:

$$\texttt{CtrlPts2Slopes} = \begin{pmatrix} 1 & 0 & 0 \\ -\Delta_{12}^{-1} & \Delta_{12}^{-1} & 0 \\ 0 & -\Delta_{23}^{-1} & \Delta_{23}^{-1} \end{pmatrix}, \tag{10}$$

where $\Delta_{ij}$ is the chromosomal position (horizontal value) of the $j$-th control point minus the position of the $i$-th control point. We can thus relate the vector of slopes $\vec{\alpha}$ and the vector of (vertical) control point values $\vec{\theta}$:

$$\vec{\alpha} = \texttt{CtrlPts2Slopes} \cdot \vec{\theta} = \begin{pmatrix} \theta_1 \\ (\theta_2 - \theta_1)/\Delta_{12} \\ (\theta_3 - \theta_2)/\Delta_{23} \end{pmatrix} \tag{11}$$

From the slopes, we can obtain the fork velocities with the following matrix:

$$\texttt{Slopes2Vels} = \begin{pmatrix} 1 & 0 & 0 \\ 0 & -k_G/\alpha_2^2 & 0 \\ 0 & 0 & -k_G/\alpha_3^2 \end{pmatrix}. \tag{12}$$

The velocities $\vec{v}$ are related to $\vec{\alpha}$ like so:

$$\vec{v} = \texttt{Slopes2Vels} \cdot \vec{\alpha} = \begin{pmatrix} \alpha_1 \\ -k_G/\alpha_2 \\ -k_G/\alpha_3 \end{pmatrix}. \tag{13}$$

To properly transform the covariance matrix obtained in Sec. III C 7, we must use the transformation matrix on both sides, like so:

$$\mathrm{cov}_\alpha = \texttt{CtrlPts2Slopes}^T \cdot \mathrm{cov}_\theta \cdot \texttt{CtrlPts2Slopes}, \tag{14}$$

$$\mathrm{cov}_v = \texttt{Slopes2Vels}^T \cdot \mathrm{cov}_\alpha \cdot \texttt{Slopes2Vels}. \tag{15}$$

The error estimates for each parameter are the square roots of the corresponding diagonal elements.

##### 9. *Behavior near the terminus*

Due to the stochastic nature of replisome progression, replication forks do not always meet at the terminus, which corresponds to the *dif* site [1]. If one fork arrives first, it can continue past the *dif* site, converting from antegrade to retrograde motion along the other arm. While there are Tus-*Ter* traps in *E. coli* and *B. subtilis* to limit the amount of retrograde motion, they are not 100% efficient and they are not present in *V. cholerae* [1]. Thus, in a population, the true termination points along the chromosome would form a distribution around the *dif site*. This means that in regions near the terminus, there can be antegrade and retrograde replication forks both contributing to the marker frequency, which contradicts one of the assumptions required for our analysis: replication proceeding only in one direction. Such retrograde motion would cause a flattening out around any fork convergence points observed in the marker-frequency data. Therefore, when doing a multi-segment analysis, we remove the two fork velocity measurements on either side of the marker-frequency minimum.

##### 10. *Selection of the number of fitted knots*

First we bin the data into 1 kb regions, following the convention of previous marker frequency analysis studies. To determine the number of segments that would best fit the data, we use the Akaike Information Criterion (AIC).

We compare a series of nested models with $2^k$ continuous piecewise-linear least squares fits, where $k = 1, ..., 10$. The AIC-optimal model for fast growth (in LB) had 39 knots, spaced by 100 kb, generating 38 measurements of locus velocity across the two chromosomes of WT *V. cholerae*. Other strains either have similar or smaller region sizes that minimized the AIC value, so for ease of comparison with other mutant strains, we take 100 kb to be the step size for determining fork velocity.

#### IV. THEORETICAL METHODS

##### A. Bilateral symmetry analysis

To analyze whether an observed velocity profile was consistent with the predictions of the time-dependent mechanism, we test for bilateral symmetry between the left and right arms:

$$v(\ell) = v(-\ell), \tag{16}$$

where $\ell$ is the locus position relative to the origin.

Assume the fork velocity has been computed over $2m$ equal-length genome segments, arranged symmetrically about the origin ($\ell = 0$). The segments $i = -1... - m$ correspond to sequentially labeled segments along the left arm and the segments $i = 1...m$ correspond to sequentially labeled segments along the right arm such that $\ell = \Delta\ell i$ corresponds to the end point of each segment where $\Delta\ell$ is the segment length. Let the mean fork velocity be defined:

$$\overline{v} \equiv \frac{1}{2m}\sum_{i=1}^{m} v_i + v_{-i}, \tag{17}$$

where $v_i$ is the velocity over segment $i$.

To divide the variance into symmetric and antisymmetric contributions, we define symmetrized, $(i)$, and antisymmetrized velocities, $[i]$, for index pairs $\pm i$:

$$\delta v_{(i)} \equiv \tfrac{1}{2}(v_i + v_{-i} - 2\overline{v}), \tag{18}$$

$$\delta v_{[i]} \equiv \tfrac{1}{2}(v_i - v_{-i}). \tag{19}$$

We define the symmetric and antisymmetric variances

$$\sigma_S^2 \equiv \frac{1}{m}\sum_{i=1}^{m} \delta v_{(i)}^2 \tag{20}$$

$$\sigma_A^2 \equiv \frac{1}{m}\sum_{i=1}^{m} \delta v_{[i]}^2, \tag{21}$$

and the total variance is the sum of the symmetric and antisymmetric variances:

$$\sigma^2 \equiv \frac{1}{2m}\sum_{i=1}^{m}(v_i - \overline{v})^2 + (v_{-i} - \overline{v})^2 \tag{22}$$

$$= \sigma_S^2 + \sigma_A^2. \tag{23}$$

Finally, we define the symmetric fraction of the variance:

$$f_S \equiv \sigma_S^2/\sigma^2. \tag{24}$$

If the variation in the velocity obeys the bilateral symmetry (Eq. 16), the variance is all symmetric (*i.e.* $f_S = 1$). If on the other hand, the variation is randomly distributed along the genome, we expect equal symmetric and antisymmetric combinations (*i.e.* $f_S = 1/2$).

##### B. Estimation of average fork number per cell cycle

The following analysis is only an estimation. There are a few key assumptions that are inconsistent with cell phenomenology, but are kept to make the analysis tractable. In particular, we take the assumption that neither arm of the chromosome has fork arrest, which is inconsistent with the results for *B. subtilis*. This assumption allows us to obtain an estimate without prior knowledge of where fork arrest may occur. We also assume that the D period (time between end of chromosomal replication and cell division) is negligible, which is inconsistent when growth is extremely slow. This assumption allows us to use the copy number of the terminus as an approximation of the number of cells, which cannot be determined from MFA.

The population number density of forks $n_f(\ell)$ at any position $\ell$ is given by the change in copy number between two consecutive positions along the chromosome:

$$n_f(\ell) = -\frac{\mathrm{d}}{\mathrm{d}\ell} N(\ell). \tag{25}$$

This equation is a result of the following reasoning: If there are two copies at $\ell = x$ and one copy at $\ell = x + 1$, then there must be a fork that has just replicated $\ell = x$. We need to integrate the number density over the entire chromosome to get the total number of forks for the population:

$$N_{fork,population} = 2\int_{ori}^{ter} n_f(\ell)\,\mathrm{d}\ell = 2\int_{ori}^{ter} -\frac{\mathrm{d}}{\mathrm{d}\ell} N(\ell)\,\mathrm{d}\ell = 2(N_{ori} - N_{ter}), \tag{26}$$

where the factor of 2 came from having two arms. To get the average number of forks per cell $\bar{N}_f$, we divide by the total number of cells, which is roughly equal to the number of termini $N_{ter}$:

$$\bar{N}_f = 2\frac{N_{ori} - N_{ter}}{N_{ter}}. \tag{27}$$

This can be related to lag time:

$$\tau_\ell = -k_G^{-1} \ln \frac{N_\ell}{N_{ori}} \implies \frac{N_{ori}}{N_{ter}} = \exp(k_G \tau_{ter}) = e^{\tau_{ter} \ln 2/T} = \left(e^{\ln 2}\right)^{\tau_{ter}/T} = 2^{\tau_{ter}/T}, \tag{28}$$

which can be substituted into the equation above to get:

$$\bar{N}_f = 2 * (2^{\tau_{ter}/T} - 1). \tag{29}$$

This is closely related to the result derived in [7]. For two chromosomes, we add the number of forks for both, but divide everything just by the minimum $N_{ter,i}$, which corresponds to the terminus of Chr1 in this case:

$$\bar{N}_{f,2chr} = 2\frac{(N_{ori,1} - N_{ter,1})}{N_{ter,1}} + 2\frac{(N_{ori,2} - N_{ter,2})}{N_{ter,1}}. \tag{30}$$

In general, when there are multiple chromosomes:

$$\bar{N}_{fork,multichr} = \frac{2}{\min(N_{ter,i})} \sum_i (N_{ori,i} - N_{ter,i}), \tag{31}$$

where $i$ denotes which chromosome. To get a relative lag time, we use:

$$\tau_{\ell \to m} = \tau_m - \tau_\ell = -k_G^{-1}\left(\ln \frac{N_m}{N_{ori}} - \ln \frac{N_\ell}{N_{ori}}\right) = k_G^{-1} \ln \frac{N_\ell}{N_m}. \tag{32}$$

Letting the subscript *end* denote the lag time corresponding to $\min(N_{ter,i})$. We thus have:

$$\bar{N}_{fork,multichr} = 2\sum_i \left(2^{(\tau_{end} - \tau_{ori,i})/T} - 2^{(\tau_{end} - \tau_{ter,i})/T}\right), \tag{33}$$

where $\tau_{end}$ is the maximum lag time measured. This can be simplified to:

$$\bar{N}_{fork,multichr} = 2 \times 2^{\tau_{end}/T} \times \sum_i \left(2^{-\tau_{ori,i}/T} - 2^{-\tau_{ter,i}/T}\right), \tag{34}$$

which enables a calculation of average fork number per cell cycle using lag times.

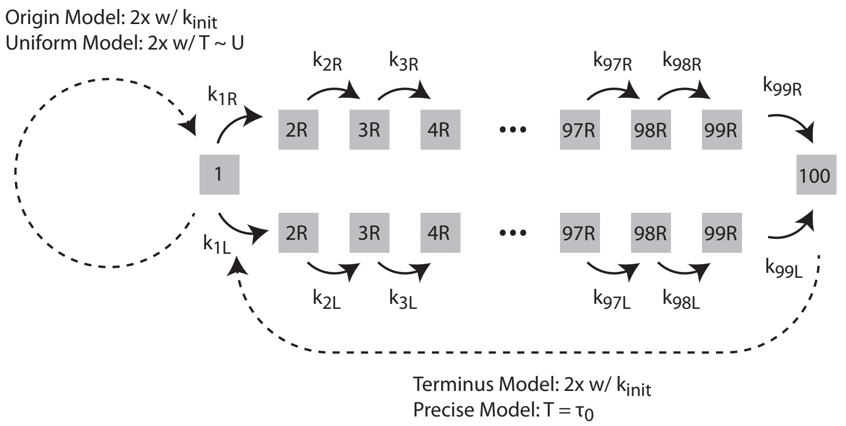


FIG. 2. **Stochastic simulation model schematics.** Replication initiates at the origin (state 1) and proceeds bi-directionally along the left (L) and right arms (R). Each transition between simulation bases (*i.e.* state number) represents the replication of roughly 25 kb. States have exponentially-distributed lifetimes with a rate equal to the fork velocity at that position: $k_X = v(\ell_X)$. A number of different initiation schemes were simulated. In the *Origin Model*, the initiation rate was proportional to the number origins (per cell). In the *Terminus Model*, the initiation rate was proportional to the number termini (per cell). In the *Uniform Model*, there is a uniformly distributed wait time after initiation before the next initiation occurs. In the *Precise Model*, there is a fixed time interval after termination before the next initiation occurs.

##### C. Statistically significant deviations of local fork velocity from the global mean

To test the statistical significance of the measured velocities, we use a null hypothesis test. The null hypothesis corresponds to a model with uniform fork velocity on the right and left arms. The alternative hypothesis corresponds to a model with a model-selected number of knots ($m$), corresponding to $m - 1$ genomic region-specific velocities.

To calculate the p-value, we begin by obtaining the $\chi^2$-test statistic:

$$\chi^2 = \sum_i \frac{(O_i - E_i)^2}{\sigma_i^2} \tag{35}$$

where $O_i$ denotes the observed values, $E_i$ denotes the expected values, and $\sigma_i^2$ is the variance of the fork velocities measured using the fitter function (as described in Supplemental Material Sec. III C 8). In this case, $E_i$ is the global mean fork velocity for all $i$, since that is the null hypothesis. Since there is only one degree of freedom for the model, we use the one-dimensional $\chi^2$ cumulative distribution function to determine the probability of measuring a $\chi^2$ statistic as extreme as the one we've measured, assuming the null hypothesis is true. This p-value is found to be $\ll 10^{-30}$ for all datasets except one with a p-value of $6 \times 10^{-12}$. All of the p-values are much smaller than the standard threshold of 0.05, so we reject the null hypothesis. We thus conclude that there are statistically significant variations of the fork velocity relative to the mean.

#### V. SUPPLEMENTAL THEORETICAL RESULTS

##### A. Stochastic simulations

###### 1. *Stochastic simulations method*

The purpose of the stochastic simulation was *to investigate the role of stochasticity in determining the log-phase growth demography of the population*. It was *not* to generate a mechanistically realistic and detailed model of the cell cycle.

To simulate the cell cycle, we performed an exact stochastic simulation using the Gillespie Algorithm [8]. We idealized the cell cycle as follows: **Genome representation:** The genome was divided into two equal length arms each consisting to 100 course-grained bases. An explicit realization of all 5 Mb would be too slow to rapidly explore different cell models. Finer coarse-grained basepairs were explored but made no difference to the results. **Replication initiation**: We experimented with a number of different models for initiation since we initially believed that the stochasticity of initiation would limit the ability to quantitate the fork velocity. (i) We initially investigate a *Terminus*

*Model* in which there was a constant rate of initiation $k_{\rm init}$ after the termination of the on-going replication. (ii) In order to simulate a more realistic model, we then considered an *Origin Model* where there was constant rate $k_{\rm init}$ of initiation at each origin. (iii) We explored a *Precise Model* with precise timed B period length $T_B$. (iv) We explored a *Uniform Model* with a uniform distribution of B periods. **Fork velocity:** We used the replication of the course-grained bases had exponentially distributed wait times $k_{\rm rep} = v/\Delta\ell$ (*i.e.* constant rate per unit time) where $v$ is the replication velocity and $\Delta\ell$ is the length of the course-grained bases. We note that this assumption makes replication *more stochastic* than a more realistic model in which wait times are exponentially distributed at the single-base level. We experimented with a number of different types of locus-dependent fork velocities. In addition, we added exponentially-distributed pauses of various durations at a specific locus. **Termination:** We only allowed a single direction of fork propagation. When the fork reached the end of the arm, replication terminated. **Cell division:** We assumed cells divided immediately after termination of the slowest arm of the chromosome. **Initiation of the simulation:** All simulations initiated from a single new-born cell with an un-replicated chromosome. **Stop conditions:** Simulation were run until there were $10^5$ cells. **Determination of growth rate.** The number of cells was fit to an exponential to determine the growth rate $k_G$ between $N_{\rm cell}(t) = 10^4$ and $N_{\rm cell}(t) = 10^5$ cells. **Generation of simulated marker frequency.** The marker frequency was defined as the number of each genetic locus in the population at the termination of the simulation. In summary, the model was described by the parameters $k_{\rm init}$ and the fork velocity $v(\ell)$ (or pause time distribution) at each locus. The model output was the marker frequency at each locus $N(\ell)$ and the growth rate $k_G$. See Fig. 2.

##### 2. *Stochastic simulations match the predictions of the log-slope*

To test whether the log-slope law applies locally, irrespective of stochasticity in cell-cycle timing, we used a stochastic simulation to generate simulated marker-frequency data. The basic strategy was to simulate a stochastically timed cell cycle, including stochastically timed B periods as well as stochastic fork dynamics, and compare the observed population demographics to the prediction of non-stochastic models.

**Log slope.** We define the log slope:

$$\alpha(\ell) \equiv \frac{\rm d}{{\rm d}\ell} \log N(\ell), \tag{36}$$

where $N(\ell)$ is the population copy number of the locus at position $\ell$. Theoretically, the slope is predicted to be:

$$\alpha(\ell) = \frac{k_G}{v(\ell)}, \tag{37}$$

where $k_G$ is the growth rate and $v(\ell)$ is the locus-dependent fork velocity.

**Log difference.** For long pauses, the dynamics are characterized by a step rather than a slope. The step is defined:

$$\Delta(\ell) \equiv \lim_{\delta\to 0^+} \log N(\ell+\delta) - \log N(\ell), \tag{38}$$

where $N(\ell)$ is the population copy number of the locus at position $\ell$. Theoretically, the log difference is predicted to be:

$$\Delta(\ell) \equiv k_G \delta\tau(\ell) = \log\big(1 + \tfrac{k_G}{k}\big), \tag{39}$$

where $k_G$ is the growth rate, $\delta\tau(\ell)$ is the exponential mean of the step wait time and the last equality applies in the special case of an exponentially-distributed wait time with rate $k$.

##### 3. *Simulation models*

The model descriptions and motivations are summarized below:

**Terminus Model 1:** In the terminus model, the replication initiation rate is proportional to the terminus number (per cell). In this model, there is a locus-independent fork velocity.

**Terminus Model 2:** Same as above, but with two fork velocities. On the last half of the right arm (region 1), the fork velocity is higher than the rest of the chromosome (region 0).

**Terminus Model 3:** Same as above, but with an exponentially-distributed pause close to the origin on the right arm.

**Terminus Model 4:** Same as Terminus 1, but with a lower initiation rate.

**Origin Model:** In the origin model, the replication initiation rate is proportional to the origin number.

**Uniform Model:** The timing between initiations is uniformly distributed. This model was included to explore whether the observations were a special case of exponentially-distributed wait times.

**Precise Model:** The timing between initiations is uniformly distributed. This model was included to explore whether the observations were a special case of exponentially-distributed wait times.

##### 4. *Simulation results*

The results from the simulations are summarized in the table below. All numbers have at least precision of 1% and are in simulation units: simulation bases (sb) and simulation time (st).

| Model | Parameter Initiation | Parameter Fork dynamics | Observed Growth rate: $k_G$ | Observed Log slope/difference | Predicted Log slope/difference |
|---|---|---|---|---|---|
| Terminus 1 | $k_{\text{init}} = 1$ st$^{-1}$ | $v_0 = 20$ sb/st | $2.6 \times 10^{-1}$ st$^{-1}$ | $\alpha = 9.0 \times 10^{-3}$ sb$^{-1}$ | $\alpha = 9.0 \times 10^{-3}$ sb$^{-1}$ |
| Terminus 2 | $k_{\text{init}} = 1$ st$^{-1}$ | $v_0 = 20$ sb/st<br>$v_1 = 30$ sb/st | $2.7 \times 10^{-1}$ st$^{-1}$ | $\alpha_0 = 1.3 \times 10^{-2}$ sb$^{-1}$<br>$\alpha_1 = 8.8 \times 10^{-3}$ sb$^{-1}$ | $\alpha_0 = 1.3 \times 10^{-2}$ sb$^{-1}$<br>$\alpha_1 = 8.8 \times 10^{-3}$ sb$^{-1}$ |
| Terminus 3 | $k_{\text{init}} = 1$ st$^{-1}$ | $v_0 = 20$ sb/st<br>$v_1 = 30$ sb/st<br>$k_3 = 2$ st$^{-1}$ | $2.6 \times 10^{-1}$ st$^{-1}$ | $\alpha_0 = 1.3 \times 10^{-2}$ sb$^{-1}$<br>$\alpha_1 = 8.8 \times 10^{-3}$ sb$^{-1}$<br>$\Delta = 1.3 \times 10^{-1}$ | $\alpha_0 = 1.3 \times 10^{-2}$ sb$^{-1}$<br>$\alpha_1 = 8.8 \times 10^{-3}$ sb$^{-1}$<br>$\Delta = 1.3 \times 10^{-1}$ |
| Terminus 4 | $k_{\text{init}} = 0.3$ st$^{-1}$ | $v_0 = 20$ sb/st | $1.4 \times 10^{-1}$ st$^{-1}$ | $\alpha_0 = 7.3 \times 10^{-3}$ sb$^{-1}$ | $\alpha_0 = 7.3 \times 10^{-3}$ sb$^{-1}$ |
| Origin | $k_{\text{init}} = 0.3$ st$^{-1}$ | $v_0 = 20$ sb/st | $3.0 \times 10^{-1}$ st$^{-1}$ | $\alpha = 1.5 \times 10^{-2}$ sb$^{-1}$ | $\alpha = 1.5 \times 10^{-2}$ sb$^{-1}$ |
| Uniform | $T \sim U([0,6])$ st | $v_0 = 20$ sb/st | $2.7 \times 10^{-1}$ st$^{-1}$ | $\alpha = 1.3 \times 10^{-2}$ sb$^{-1}$ | $\alpha = 1.3 \times 10^{-2}$ sb$^{-1}$ |
| Precise | $T = 0.2$ st | $v_0 = 20$ sb/st | $1.4 \times 10^{-1}$ st$^{-1}$ | $\alpha = 6.8 \times 10^{-3}$ sb$^{-1}$ | $\alpha = 6.8 \times 10^{-3}$ sb$^{-1}$ |

##### 5. *Re-scaling simulation units to compare to measured data*

The simulations were performed in simulation units. We call the coarse-grained bases simulation bases (sb) and time units simulation time (st). To compare these simulations to experimental data, the prediction need to be rescaled to place the observations in biological units. To model a 5 Mb bacterial genome, with typical fork velocity of 1 kb/s, the conversion factors are:

$$\frac{5 \times 10^6 \text{ bp}}{200 \text{ sb}} = 2.5 \times 10^4 \ \frac{\text{bp}}{\text{sb}}, \tag{40}$$

$$\left(\frac{1 \text{ s}}{10^3 \text{ bp}}\right)\left(\frac{20 \text{ sb}}{1 \text{ st}}\right)\left(\frac{5 \times 10^6 \text{ bp}}{200 \text{ sb}}\right) = 5 \times 10^2 \ \frac{\text{s}}{\text{st}}, \tag{41}$$

since we simulated typical fork velocities of 20 sb/st which we rescale the units to be equivalent to 1 kb/s.

**Aside:** Note that due to the smaller number of sb relative to bp, the stochasticity of fork dynamics should be exaggerated in the simulations, strengthening our argument that the stochasticity does not effect the model predictions.

##### 6. *Movies of marker frequency dynamics approaching steady state growth*

We include two examples of movies tracking the marker frequency dynamics from a single cell to steady state growth. Movies `term1_mov.mp4` and `term3_mov.mp4` show the dynamics of the Terminus 1 and Terminus 3 models respectively. Movie time is shown in simulation units. The dynamics emphasizes the importance of performing the calculations of the marker frequency in the steady-state limit. Not taking this limit correctly leads to anomalous results (*e.g.* [9]).

### VI. SUPPLEMENTAL EXPERIMENTAL RESULTS

#### A. Comparison with GC content

One potential rate-limiting factor for replication is the distribution of GC content across the genome. However, at the current genomic resolution of 100 kb, we find no significant evidence of GC content determining the rate of replication. See Fig. 3 for the results of wild-type *E. coli* strain MG1655 in LB and M9.

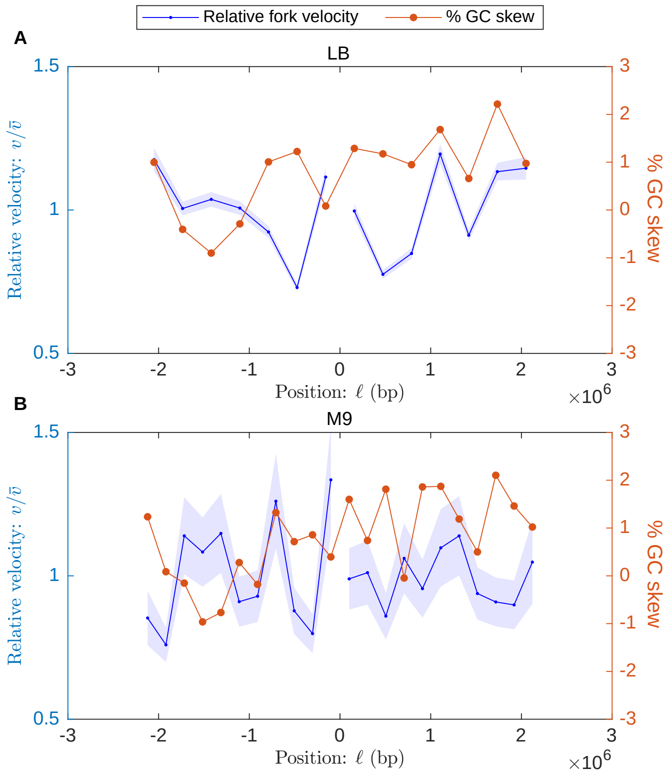


FIG. 3. **Relative fork velocity and % GC skew as a function of position.** Results for wild-type *E. coli* strain MG1655. The blue curve represents the fork velocity divided by the mean fork velocity, with shaded error bars. The orange curve represents the % GC skew. **Panel A:** Growth in LB medium. **Panel B:** Growth in M9 glucose minimal medium.

#### B. *V. cholerae* WT on LB

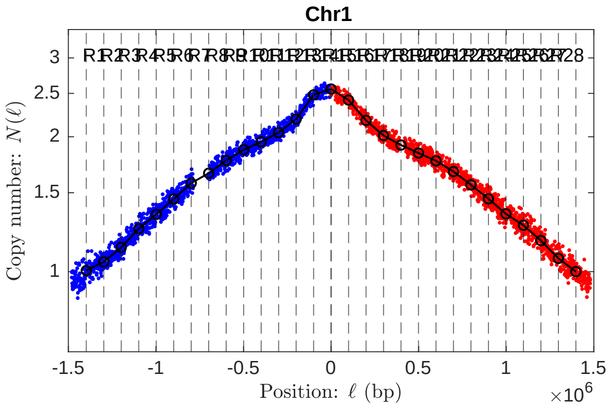


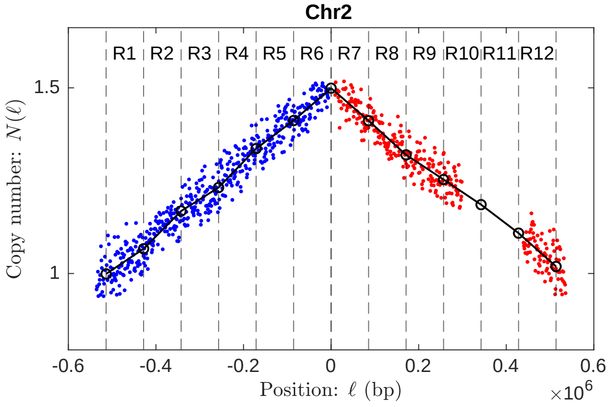


Detailed results tabulated in `SuppData.xlsx`.

##### C. *V. cholerae* WT on M9 fructose

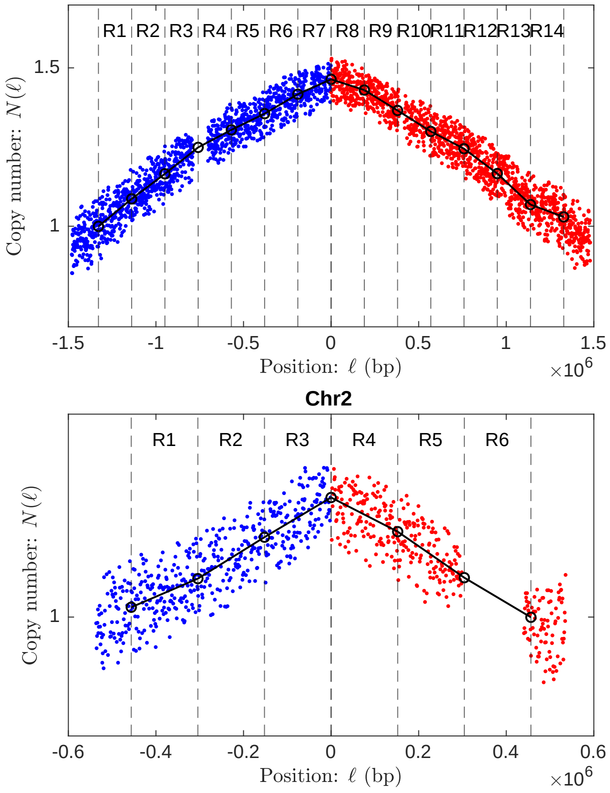


Detailed results tabulated in `SuppData.xlsx`.

##### D. *V. cholerae* MCH1 on LB

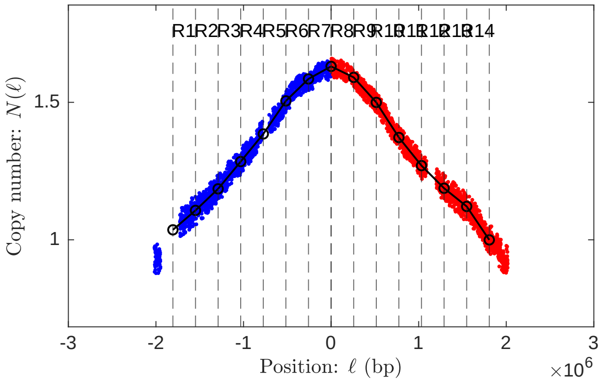


Detailed results tabulated in `SuppData.xlsx`.

##### E. *V. cholerae* MCH1 on M9 fructose

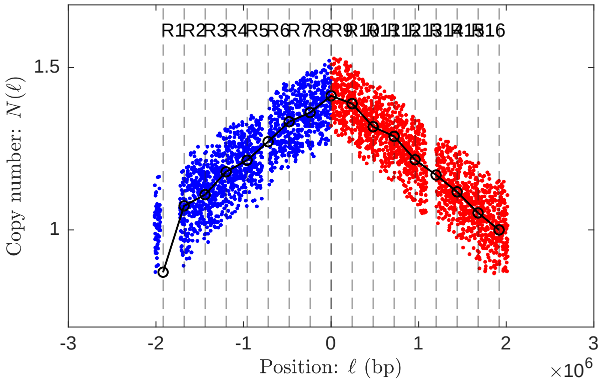


Detailed results tabulated in `SuppData.xlsx`.

##### F. *V. cholerae oriR4* on M9 fructose

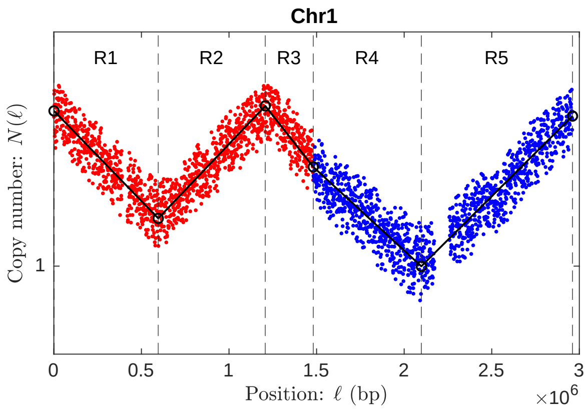


| Region name | Region start position: $\ell_-$ (Mb) | Region end position: $\ell_+$ (Mb) | Fork direction | Replication orientation | Slope: $\alpha$ (Mb$^{-1}$) | Velocity: $v$ (kb/s) | Log difference: $\Delta$ | Duration: $\Delta\tau$ (min) | Percent error |
|---|---|---|---|---|---|---|---|---|---|
| R1 | 0 | 0.596 | + | A | 0.308 | 0.750 | 0.183 | 13.2 | ±2.0% |
| R2 | 0.596 | 1.21 | - | R | 0.314 | 0.736 | 0.192 | 13.8 | ±1.7% |
| R3 | 1.21 | 1.48 | + | A | 0.380 | 0.608 | 0.104 | 7.51 | ±3.3% |
| R4 | 1.48 | 2.10 | + | R | 0.275 | 0.840 | 0.170 | 12.2 | ±1.9% |
| R5 | 2.10 | 2.96 | - | A | 0.297 | 0.777 | 0.256 | 18.5 | ±1.3% |

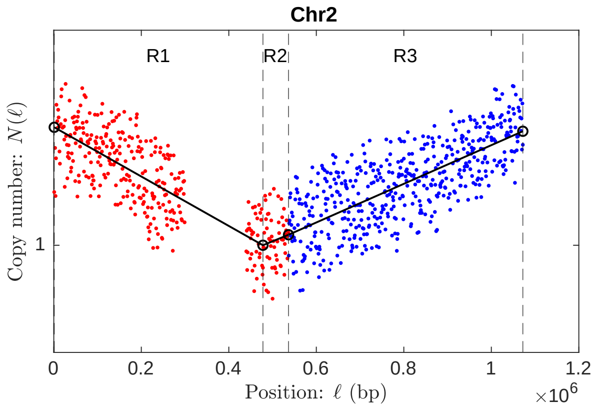


| Region name | Region start position: $\ell_-$ (Mb) | Region end position: $\ell_+$ (Mb) | Fork direction | Replication orientation | Slope: $\alpha$ (Mb$^{-1}$) | Velocity: $v$ (kb/s) | Log difference: $\Delta$ | Duration: $\Delta\tau$ (min) | Percent error |
|---|---|---|---|---|---|---|---|---|---|
| R1 | 0 | 0.478 | + | A | 0.257 | 0.898 | 0.123 | 8.85 | ±5.0% |
| R2 | 0.478 | 0.537 | - | R | 0.186 | 1.24 | 0.0109 | 0.787 | ±52% |
| R3 | 0.537 | 1.07 | - | A | 0.201 | 1.15 | 0.108 | 7.76 | ±4.4% |

##### G. *B. subtilis oriC*-257° on S7 fumarate

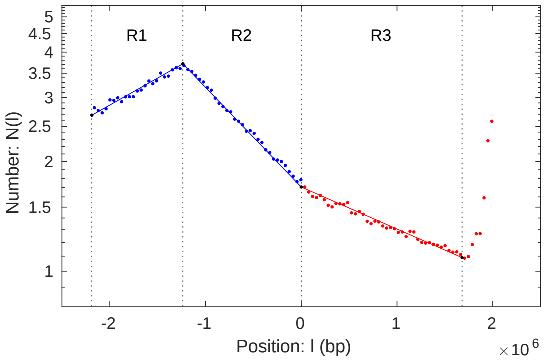


| Region name | Region start position: $\ell_-$ (Mb) | Region end position: $\ell_+$ (Mb) | Fork direction | Replication orientation | Slope: $\alpha$ (Mb$^{-1}$) | Velocity: $v$ (kb/s) | Log difference: $\Delta$ | Duration: $\Delta\tau$ (min) |
|---|---|---|---|---|---|---|---|---|
| R1 | $-2.19$ | $-1.24$ | - | A | $0.342 \pm 4.3\%$ | NA | $0.325 \pm 4.3\%$ | NA |
| R2 | $-1.24$ | $-8.72 \times 10^{-4}$ | + | R | $0.631 \pm 1.3\%$ | NA | $0.779 \pm 1.3\%$ | NA |
| R3 | $-8.72 \times 10^{-4}$ | 1.68 | + | R | $0.266 \pm 2.6\%$ | NA | $0.447 \pm 2.6\%$ | NA |

##### H. *B. subtilis oriC*-94° on S7 fumarate

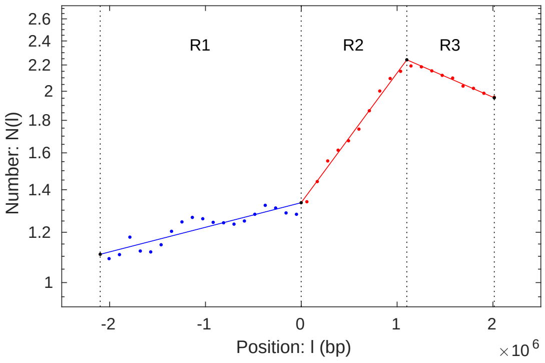


| Region name | Region start position: $\ell_-$ (Mb) | Region end position: $\ell_+$ (Mb) | Fork direction | Replication orientation | Slope: $\alpha$ (Mb$^{-1}$) | Velocity: $v$ (kb/s) | Log difference: $\Delta$ | Duration: $\Delta\tau$ (min) |
|---|---|---|---|---|---|---|---|---|
| R1 | $-2.10$ | $-1.44 \times 10^{-3}$ | - | A | $8.91 \times 10^{-2} \pm 10\%$ | NA | $0.187 \pm 10\%$ | NA |
| R2 | $-1.44 \times 10^{-3}$ | 1.10 | - | R | $0.469 \pm 3.5\%$ | NA | $0.518 \pm 3.5\%$ | NA |
| R3 | 1.10 | 2.01 | + | A | $0.151 \pm 18\%$ | NA | $0.138 \pm 18\%$ | NA |

#### I. *B. subtilis oriN* on S7 fumarate

##### 1. *Two-slopes model*

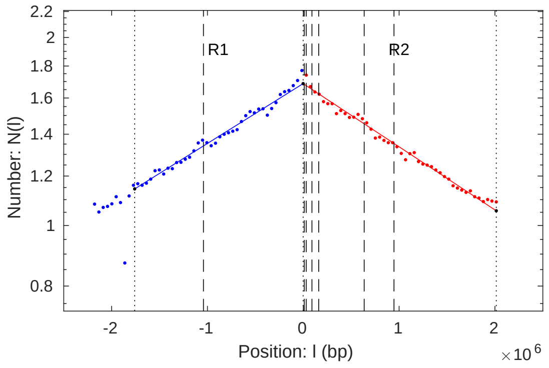


| Region name | Region start position: $\ell_-$ (Mb) | Region end position: $\ell_+$ (Mb) | Fork direction | Replication orientation | Slope: $\alpha$ (Mb$^{-1}$) | Velocity: $v$ (kb/s) | Log difference: $\Delta$ | Duration: $\Delta\tau$ (min) |
|---|---|---|---|---|---|---|---|---|
| R1 | $-1.76$ | $-1.59 \times 10^{-3}$ | - | A | $0.221 \pm 2.3\%$ | $0.948 \pm 2.3\%$ | $0.388 \pm 2.3\%$ | $30.9 \pm 2.3\%$ |
| R2 | $-1.59 \times 10^{-3}$ | 2.01 | + | A | $0.232 \pm 1.8\%$ | $0.900 \pm 1.8\%$ | $0.469 \pm 1.8\%$ | $37.3 \pm 1.8\%$ |

##### 2. *Pause model*

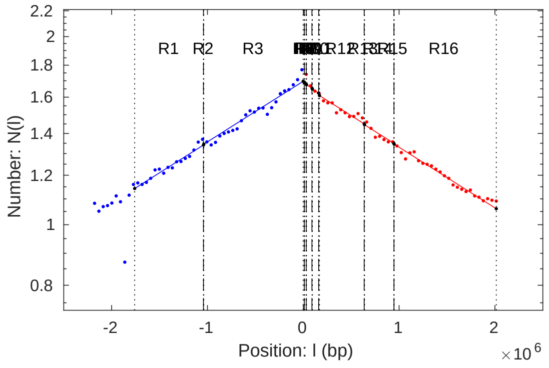


| Region name | Region start position: $\ell_-$ (Mb) | Region end position: $\ell_+$ (Mb) | Fork direction | Replication orientation | Slope: $\alpha$ (Mb$^{-1}$) | Velocity: $v$ (kb/s) | Log difference: $\Delta$ | Duration: $\Delta\tau$ (min) |
|---|---|---|---|---|---|---|---|---|
| R1 | $-1.76$ | $-1.04$ | - | A | $0.223 \pm 1.9\%$ | $0.937 \pm 1.9\%$ | $0.160 \pm 1.9\%$ | $12.8 \pm 1.9\%$ |
| R2 | $-1.04$ | $-1.04$ | - | A | $0.973 \pm 21\%$ | $0.215 \pm 21\%$ | $3.13 \times 10^{-3} \pm 21\%$ | $0.250 \pm 21\%$ |
| R3 | $-1.04$ | $-1.59 \times 10^{-3}$ | - | A | $0.223 \pm 1.9\%$ | $0.937 \pm 1.9\%$ | $0.232 \pm 1.9\%$ | $18.4 \pm 1.9\%$ |
| R4 | $-1.59 \times 10^{-3}$ | $1.13 \times 10^{-2}$ | + | R | $0.223 \pm 1.9\%$ | $0.937 \pm 1.9\%$ | $2.88 \times 10^{-3} \pm 1.9\%$ | $0.229 \pm 1.9\%$ |
| R5 | $1.13 \times 10^{-2}$ | $1.45 \times 10^{-2}$ | + | A | $0.973 \pm 21\%$ | $0.215 \pm 21\%$ | $3.13 \times 10^{-3} \pm 21\%$ | $0.250 \pm 21\%$ |
| R6 | $1.45 \times 10^{-2}$ | $3.06 \times 10^{-2}$ | + | A | $0.223 \pm 1.9\%$ | $0.937 \pm 1.9\%$ | $3.59 \times 10^{-3} \pm 1.9\%$ | $0.286 \pm 1.9\%$ |
| R7 | $3.06 \times 10^{-2}$ | $3.38 \times 10^{-2}$ | + | A | $0.973 \pm 21\%$ | $0.215 \pm 21\%$ | $3.13 \times 10^{-3} \pm 21\%$ | $0.250 \pm 21\%$ |
| R8 | $3.38 \times 10^{-2}$ | $9.18 \times 10^{-2}$ | + | A | $0.223 \pm 1.9\%$ | $0.937 \pm 1.9\%$ | $1.29 \times 10^{-2} \pm 1.9\%$ | $1.03 \pm 1.9\%$ |
| R9 | $9.18 \times 10^{-2}$ | $9.50 \times 10^{-2}$ | + | A | $0.973 \pm 21\%$ | $0.215 \pm 21\%$ | $3.13 \times 10^{-3} \pm 21\%$ | $0.250 \pm 21\%$ |
| R10 | $9.50 \times 10^{-2}$ | 0.159 | + | A | $0.223 \pm 1.9\%$ | $0.937 \pm 1.9\%$ | $1.44 \times 10^{-2} \pm 1.9\%$ | $1.15 \pm 1.9\%$ |
| R11 | 0.159 | 0.169 | + | A | $0.973 \pm 21\%$ | $0.215 \pm 21\%$ | $9.40 \times 10^{-3} \pm 21\%$ | $0.749 \pm 21\%$ |
| R12 | 0.169 | 0.636 | + | A | $0.223 \pm 1.9\%$ | $0.937 \pm 1.9\%$ | $0.104 \pm 1.9\%$ | $8.30 \pm 1.9\%$ |
| R13 | 0.636 | 0.639 | + | A | $0.973 \pm 21\%$ | $0.215 \pm 21\%$ | $3.13 \times 10^{-3} \pm 21\%$ | $0.250 \pm 21\%$ |
| R14 | 0.639 | 0.945 | + | A | $0.223 \pm 1.9\%$ | $0.937 \pm 1.9\%$ | $6.83 \times 10^{-2} \pm 1.9\%$ | $5.44 \pm 1.9\%$ |
| R15 | 0.945 | 0.948 | + | A | $0.973 \pm 21\%$ | $0.215 \pm 21\%$ | $3.13 \times 10^{-3} \pm 21\%$ | $0.250 \pm 21\%$ |
| R16 | 0.948 | 2.01 | + | A | $0.223 \pm 1.9\%$ | $0.937 \pm 1.9\%$ | $0.238 \pm 1.9\%$ | $19.0 \pm 1.9\%$ |

##### J. *B. subtilis oriN*-257° on S7 fumarate

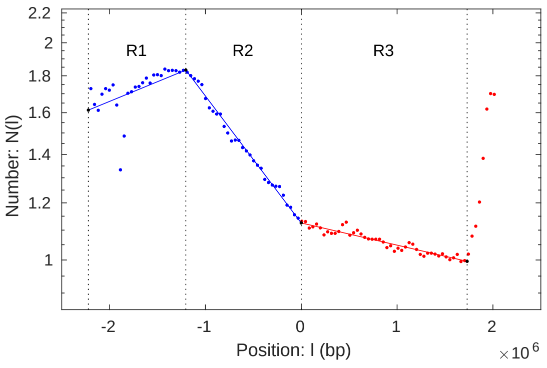


| Region name | Region start position: $\ell_-$ (Mb) | Region end position: $\ell_+$ (Mb) | Fork direction | Replication orientation | Slope: $\alpha$ (Mb$^{-1}$) | Velocity: $v$ (kb/s) | Log difference: $\Delta$ | Duration: $\Delta\tau$ (min) |
|---|---|---|---|---|---|---|---|---|
| R1 | $-2.22$ | $-1.20$ | - | A | $0.126 \pm 18\%$ | NA | $0.128 \pm 18\%$ | NA |
| R2 | $-1.20$ | $-9.18 \times 10^{-4}$ | + | R | $0.405 \pm 3.5\%$ | NA | $0.488 \pm 3.5\%$ | NA |
| R3 | $-9.18 \times 10^{-4}$ | 1.73 | + | A | $7.08 \times 10^{-2} \pm 16\%$ | NA | $0.123 \pm 16\%$ | NA |

##### K. *B. subtilis rrnIHG* inversion on MOPS glucose w/ Casamino Acids

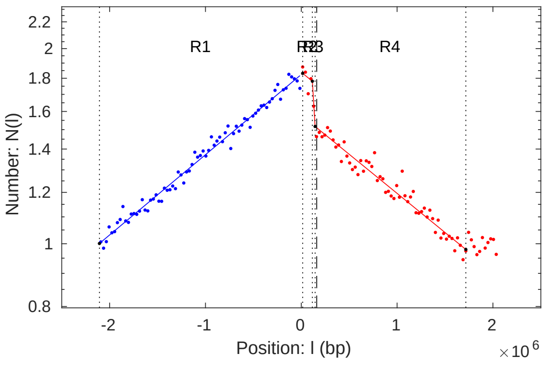


| Region name | Region start position: $\ell_-$ (Mb) | Region end position: $\ell_+$ (Mb) | Fork direction | Replication orientation | Slope: $\alpha$ (Mb$^{-1}$) | Velocity: $v$ (kb/s) | Log difference: $\Delta$ | Duration: $\Delta\tau$ (min) |
|---|---|---|---|---|---|---|---|---|
| R1 | $-2.13$ | $-6.09 \times 10^{-3}$ | - | A | $0.285 \pm 2.3\%$ | $0.921 \pm 2.3\%$ | $0.605 \pm 2.3\%$ | $38.4 \pm 2.3\%$ |
| R2 | $-6.09 \times 10^{-3}$ | $9.49 \times 10^{-2}$ | + | A | $0.278 \pm 4.0\%$ | $0.945 \pm 4.0\%$ | $2.81 \times 10^{-2} \pm 4.0\%$ | $1.78 \pm 4.0\%$ |
| R3 | $9.49 \times 10^{-2}$ | 0.124 | + | A | $5.57 \pm 8.4\%$ | $4.71 \times 10^{-2} \pm 8.4\%$ | $0.161 \pm 8.4\%$ | $10.2 \pm 8.4\%$ |
| R4 | 0.124 | 1.70 | + | A | $0.278 \pm 4.0\%$ | $0.945 \pm 4.0\%$ | $0.437 \pm 4.0\%$ | $27.7 \pm 4.0\%$ |

##### L. *B. subtilis rrnIHG* inversion on MOPS glucose – Minimal

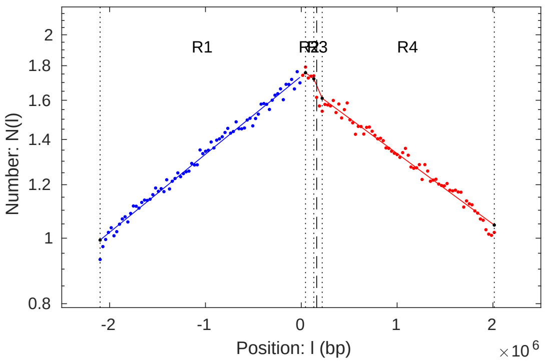


| Region name | Region start position: $\ell_-$ (Mb) | Region end position: $\ell_+$ (Mb) | Fork direction | Replication orientation | Slope: $\alpha$ (Mb$^{-1}$) | Velocity: $v$ (kb/s) | Log difference: $\Delta$ | Duration: $\Delta\tau$ (min) |
|---|---|---|---|---|---|---|---|---|
| R1 | $-2.10$ | $4.41 \times 10^{-2}$ | - | A | $0.266 \pm 1.7\%$ | $0.869 \pm 1.7\%$ | $0.570 \pm 1.7\%$ | $41.1 \pm 1.7\%$ |
| R2 | $4.41 \times 10^{-2}$ | 0.131 | + | A | $0.241 \pm 2.5\%$ | $0.961 \pm 2.5\%$ | $2.09 \times 10^{-2} \pm 2.5\%$ | $1.51 \pm 2.5\%$ |
| R3 | 0.131 | 0.218 | + | A | $0.761 \pm 13\%$ | $0.304 \pm 13\%$ | $6.61 \times 10^{-2} \pm 13\%$ | $4.77 \pm 13\%$ |
| R4 | 0.218 | 2.01 | + | A | $0.241 \pm 2.5\%$ | $0.961 \pm 2.5\%$ | $0.432 \pm 2.5\%$ | $31.2 \pm 2.5\%$ |

##### M. *E. coli* WT on LB

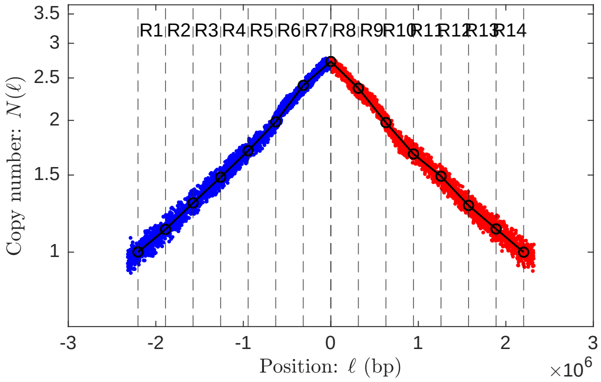


Detailed results tabulated in `SuppData.xlsx`.

##### N. *E. coli* WT on M9 glucose

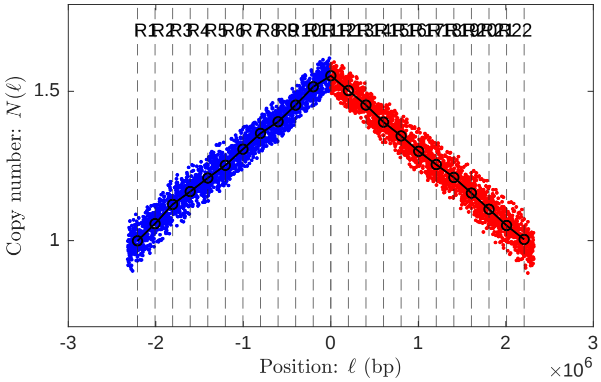


Detailed results tabulated in `SuppData.xlsx`.
